# Neuro-Metabolic and Vascular Dysregulation are Functional Biomarkers of Acquired Brain Injury

**DOI:** 10.64898/2026.10.21.760721

**Authors:** Juan Antonio K. Chong Chie, Scott A. Persohn, Amandine Jullienne, JiaMin Yan, Margaret Bright, Andre Obenaus, Paul R. Territo

## Abstract

Emerging clinical and neuropathological evidence suggests a direct link between traumatic brain injury (TBI) and neurodegenerative diseases (ND), but the mechanisms by which TBI contributes to ND onset and progression remain unclear. To bridge this gap, the NINDS TBI-ADRD consortium developed a diagnostic framework to guide the design and evaluation of TBI-ADRD experiments using translational neuroimaging (DCE-MRI, ^18^F-FDG-PET) and advanced image-processing pipelines. Neuro-metabolic and vascular dysregulation (MVD) and metabolic functional connectomics (MFC) were evaluated in 8-month C57BL/6J mice at 30- and 60-days post-injury (DPI). Both MVD and MFC revealed significant changes at 30- and 60-DPI that were sex- and injury side-dependent. Female mice showed greater regional MVD alterations, while MFC identified sex-dependent changes at 30-DPI that diminished by 60-DPI. These findings suggest that a single mid-life closed-head injury induces sexually dimorphic MVD and brain network remodeling, supporting the potential of MVD as a sensitive biomarker for improved clinical monitoring post-TBI.

## 1. INTRODUCTION

Traumatic brain injury (TBI) is a global health concern, affecting an estimated 50 to 60 million individuals annually. TBI results in long-term disabilities, including cognitive, emotional, and physical impairments, which impact both affected individuals and their caregivers.^1,2^ The associated worldwide economic burden approaches hundreds of billions of US dollars each year.^1–3^ Beyond its acute effects, this substantial burden raises important questions about whether TBI may accelerate or amplify downstream neurodegenerative processes.^4–8^ One such process is Alzheimer’s disease (AD) and related dementias (RD) which affect over 55 million people worldwide and are characterized by progressive cognitive and memory decline, typically emerging later in life, and are a leading cause of disability and dependency among the older population.^9^

Dependent upon injury severity, TBI initiates a cascade of immediate changes in the brain, beginning within minutes to days after injury.^10,11^ These changes include neuroinflammation,^12,13^ which results in a cascade of cyto- and chemokine^14^ release that reduces cerebral blood flow,^15^ resulting in continued hypometabolism,^16^ edema,^17^ and blood-brain barrier leakage.^13^ These effects may persist or worsen due to chronic neuroinflammation^12^ and increased oxidative stress,^18^ driving amyloid-beta (Aβ) and tau tangles accumulation,^19^ resulting in incomplete cerebrovascular recovery^20^ and ongoing degeneration of both white and gray matter.^21^ These pathological conditions alter the brain’s immune response,^22^ exacerbate tissue damage,^18^ and induce vascular dysfunction,^23,24^ collectively impairing cognitive and neurological function,^12,20,25^ as well as hindering proper cellular post-injury recovery.^12^ Importantly, emerging studies reveal that TBI-induced outcomes are sexually-dimorphic and are influenced by endogenous brain recovery mechanisms.^26–28^ For example, females may experience more pronounced cerebral blood flow deficits and delayed repair of vascular integrity compared to males.^29^

Emerging clinical and neuropathological evidence suggests an overlap between TBI and ADRD, proposing that TBI may accelerate AD-like pathological processes in certain individuals.^4–7^ TBI disrupts neuro-metabolic and vascular homeostasis,^16,30^ resulting in impaired cerebral perfusion,^15,20,23^ blood-brain barrier dysfunction,^13^ altered neurovascular coupling,^31^ and persistent abnormalities in glucose and mitochondrial energy metabolism.^16,23,32^ These abnormalities persist beyond the acute injury phase.^16^ Such changes may facilitate chronic tissue injury,^33^ promote Aβ and tau accumulation,^34^ impair protein clearance,^34^ and drive progressive neurodegeneration.^4,33^ Similar neurometabolic and vascular dysregulation (MVD) is observed in ADRD,^35–38^ supporting the idea that TBI may initiate or amplify pathophysiological pathways that lead to later-life neurodegeneration.^4^

Beyond MVD alterations, evidence suggests that TBI also disrupts brain neural connectivity and the organization of brain networks,^39–42^ affecting both their structure and functions.^41,43^ Disruptions in network organization,^39^ hub integrity,^44^ and inter-regional communication represent both adaptive and maladaptive response,^41,42^ altering the efficiency of information processing and coordination across brain regions.^39,40,42^ These network-level changes often accompany shifts in metabolic complexity and regional coupling,^16,45^ suggesting that TBI affects distributed brain systems rather than isolated regions.^16^ Similar network and metabolic alterations in ADRD further support the idea of shared pathophysiological mechanisms between TBI and neurodegeneration.^6,34,46^ Despite these advances, a substantial gap remains in understanding how TBI contributes to the onset and progression of neurodegenerative diseases (ND), especially regarding the temporal evolution of MVD, disturbances in brain connectivity, and their roles as drivers of neurodegeneration.

Through the NINDS PAR-23-218 *(“Development and Characterization of Experimental Models of post-TBI ADRD”*), our labs sought to develop, validate, and standardize a preclinical closed-head injury (CHI) model, and establish the construct and face validity of this model by exploring the utility of longitudinal clinically relevant translatable imaging biomarkers to characterize post-TBI pathology for alignment to clinical ADRD endophenotypes. As a first step towards this goal, we characterized CHI-driven MVD, metabolic functional connectomics (MFC) changes, and alterations in region-set enrichment analysis (RSEA)^47^ in middle-aged C57BL6/J mice using our novel imaging biomarker framework. We *hypothesized* that TBI would induce sex-specific changes in MVD and MFC, resulting in evolving regional and temporal changes in MVD phenotypes, metabolic brain networks, and RSEA outcomes. Accordingly, we aim to provide a diagnostic framework to guide the design and evaluation of TBI-ADRD, thereby advancing insight into the shared and distinct mechanisms underlying TBI and neurodegeneration prior to conducting these experiments in models of ADRD.

## 2. RESULTS

### 2.1 Neuro-Metabolic and Vascular Dysregulation Analysis

The brain regional MVD phenotypes were determined by projecting the z-scored mean values (relative to a reference) for cerebral perfusion (x-axis) and cerebral metabolism (y-axis) readouts in a 2D Cartesian coordinate system, per our published methods.^36,48^ This approach yields four possible MVD phenotypes (**Figure 1A**): Type 1 Uncoupling (↑perfusion, ↓metabolism; T1U), Type 2 Uncoupling (↓perfusion, ↑metabolism; T2U), Prodromal Coupled (↑perfusion, ↑metabolism; PC), and Neuro-Metabolic-Vascular Failure (↓perfusion, ↓metabolism; NMVF). Then, using our MVD analysis framework (**Figure S1**),^36,48^ we assessed regional metabolism and perfusion changes in the closed-head injury (CHI) model in midlife (8-months) C57BL/6J mice relative to Sham at 30-DPI and 60-DPI in each hemisphere (ipsilateral/contralateral; injured/uninjured) according to the study design (**Figure 1B**).

**Figure 1.**
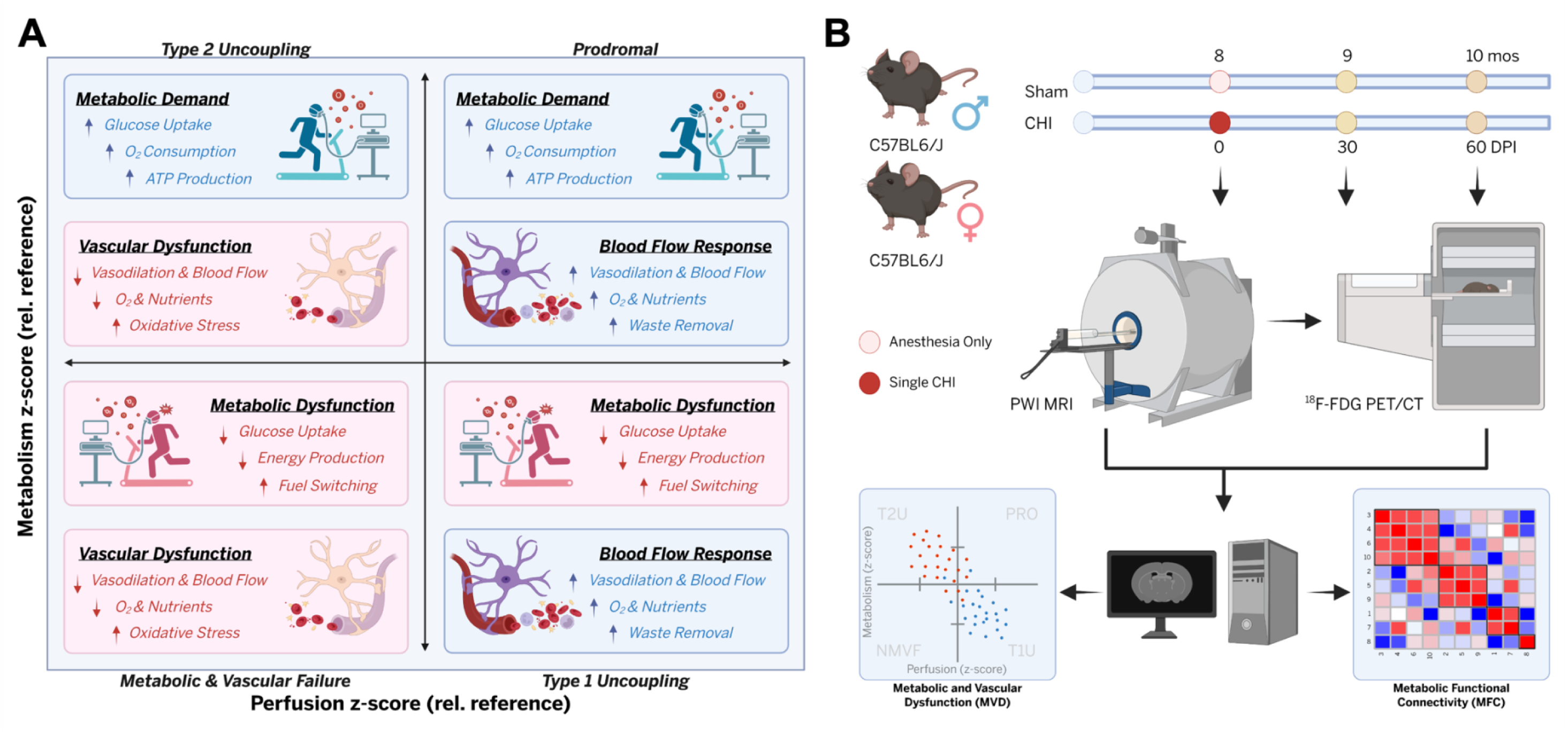
Neuro-Metabolic and Vascular Phenotypes and Study Workflow Diagram. **(A)** Conceptual 2D diagram in which the x-axis and y-axis represent the cerebral perfusion and cerebral metabolism, respectively. Each quadrant depicts one of four neuro-metabolic and vascular phenotypes, with conceptual illustrations of the corresponding metabolic and vascular/perfusion alterations. **(B)** Study workflow diagram, which describes the experimental timeline, procedures, sexes, and readouts used in this proof-of-principle study. Briefly, 8-month-old male and female mice (n=4-5/sex/treatment) received a single closed-head injury (CHI) or Sham (anesthesia only) over the left Primary (M1) and Secondary (M2) Motor cortexes, and behavior, perfusion-weighted MRI (PWI), and ^18^F-FDG PET imaging were conducted at 30 and 60 days post-injury (DPI). MVD and metabolic connectomics analyses were conducted on the 30-DPI and 60-DPI datasets.

At 30-DPI, the ipsilateral hemisphere (ILH) exhibited MVD in numerous key brain regions (e.g., Primary Motor Cortex (M1), Secondary Motor Cortex (M2), Retrosplenial Dysgranular Cortex (RSC)), which were directly impacted in CHI. For a complete list of brain regions, see **Table S1**. In male ILH of CHI relative to Sham, most regions (10 of 20; **Figure 2A**) were in the PC phenotype, whereas in females, they were predominantly in T2U (12 of 20; **Figure 2A**). A similar pattern was observed in the contralateral hemispheres (CLH): most regions in males (13 of 19; **Figure 2B**) were PC, and most regions in females (15 of 19; **Figure 2B**) were T2U. The spatial locations of regional p-values that significantly changed were projected onto the Allen MRI Template for males (**Figure 2C**) and females (**Figure 2D**), showing that altered MVD occurs not only at the impact site and neighboring regions but also on the opposite hemisphere. These findings demonstrate that MVD variations depend on sex, injury location, and affected hemisphere. For example, males tend to show increases in both cerebral perfusion and cerebral metabolism (PC phenotype), while females show decreased perfusion but increased metabolism (T2U phenotype).

**Figure 2.**
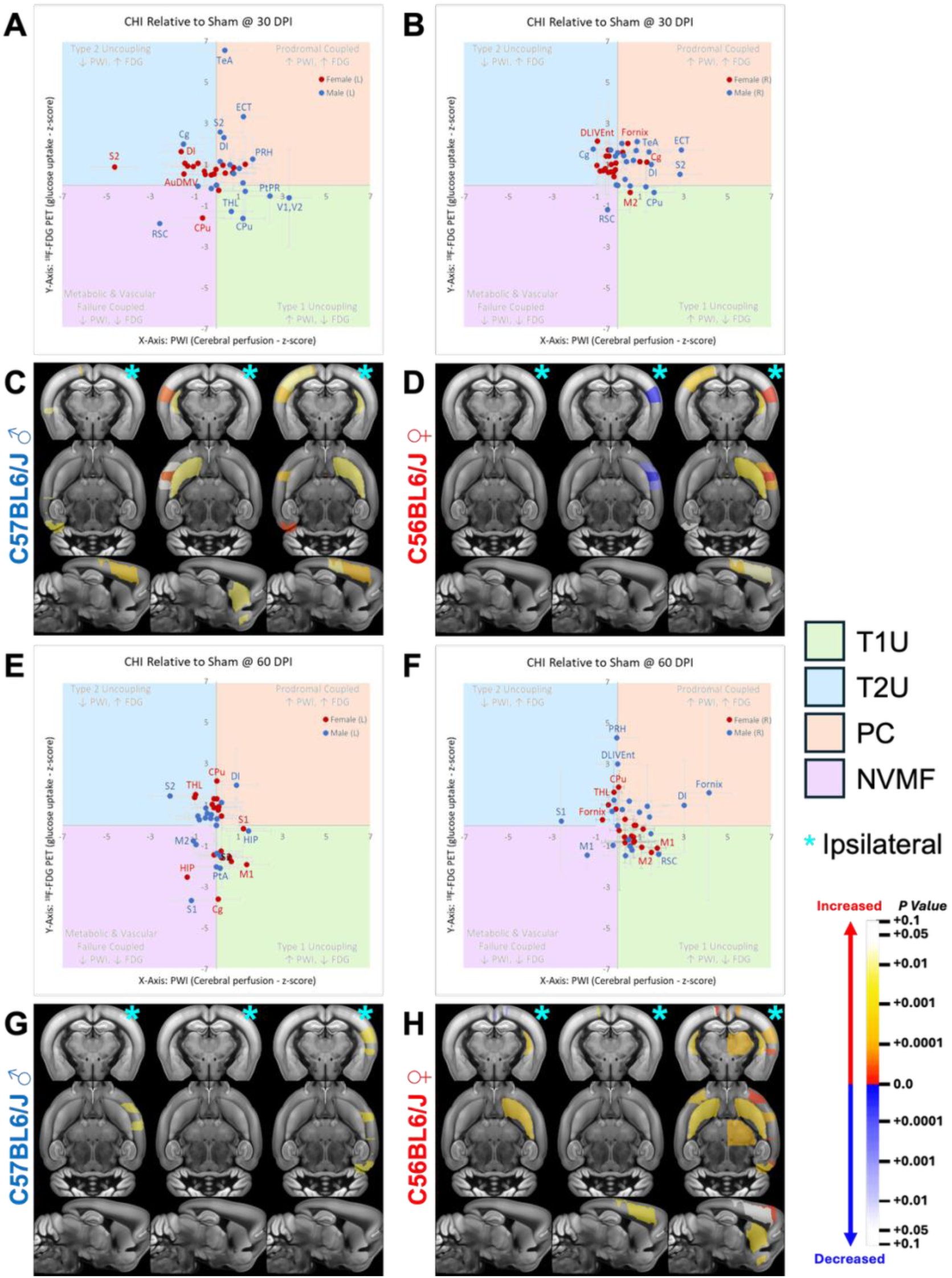
Neuro-Metabolic and Vascular Dysregulation Analysis of Ipsilateral and Contralateral Hemispheres. Neuro-Metabolic and Vascular Dysregulation (MVD) analysis of CHI relative to Sham at 30-DPI and 60-DPI for the **(A, E)** ipsilateral (ILH) and **(B, F)** contralateral (CLH) hemispheres, with signed p-value maps projected onto the Allen MRI Brain Template for **(C, G)** males and **(D, H)** females. In the 2D Cartesian MVD plots **(A, B, E, F)**, blue and red dots represent different brain regions for males and females, respectively. The cyan asterisk (*) denotes the impacted hemisphere. Significant regions (p<0.05) are labeled in the 2D MVD chart. In the p-value projections **(C, D, G, H)**, warmer colors show increased changes, while cooler colors show decreases. At both 30-DPI and 60-DPI, MVD is altered in several key regions (e.g., M1, M2, RSC) associated with the impact. Sex-dependent differences are observed at both time points and hemispheres, indicating that MVD alterations and recovery following injury depend on sex, affected hemisphere, and injury location. For a complete list of brain regions, see **Table S1**.

Analysis of the 60-DPI data revealed that altered brain regions in the ILH in both sexes were predominantly classified as T1U and T2U phenotypes (15 of 20 in males and 12 of 20 in females; **Figure 2E**). In contrast, CLH brain regions (**Figure 2F**) displayed distinct phenotypes where males exhibited T1U and PC phenotypes (13 of 19), whereas females predominantly showed T1U (14 of 19). These findings suggest that certain brain regions in males (Dorso-Lateral Intermediate Entorhinal Cortex (DLIVEnt), S2, RSC) and in females (Cingulate Cortex (Cg), M1, M2) possess sex-dependent capacities for MVD recovery following injury, indicating that the MVD is sex-dependent and temporally dynamic by brain region.

To visualize regional MVD trajectories, the 2D Cartesian charts (**Figures 2A/2B/2E/2F**) were converted into MVD migration charts (MMC; **Figure 3**), which describe the movement of brain regions across MVD phenotypes at 30- and 60-DPI. The MMC suggests that TBI has a distinct effect (i.e., larger symbol and/or greater quadrant progression) in males compared to females in brain regions such as the Temporal Association Cortex (TeA), M2, RSC, and S2. In males, comparison between both time points revealed a shift toward T1U/T2U in 8 of 20 regions in the ILH (**Figures 3A/3B**) and 7 of 19 regions in the CLH (**Figures 3C/3D**). In females, 8 of 20 for ILH (**Figures 3E/3F**) and 11 of 19 for CLH (**Figures 3G/3H**) transition towards T1U/T2U. This indicates a progressive shift toward MVD across both sexes and hemispheres. In summary, these changes likely contribute to the long-term loss of coupling mechanisms after injury in a temporally evolving manner.

**Figure 3.**
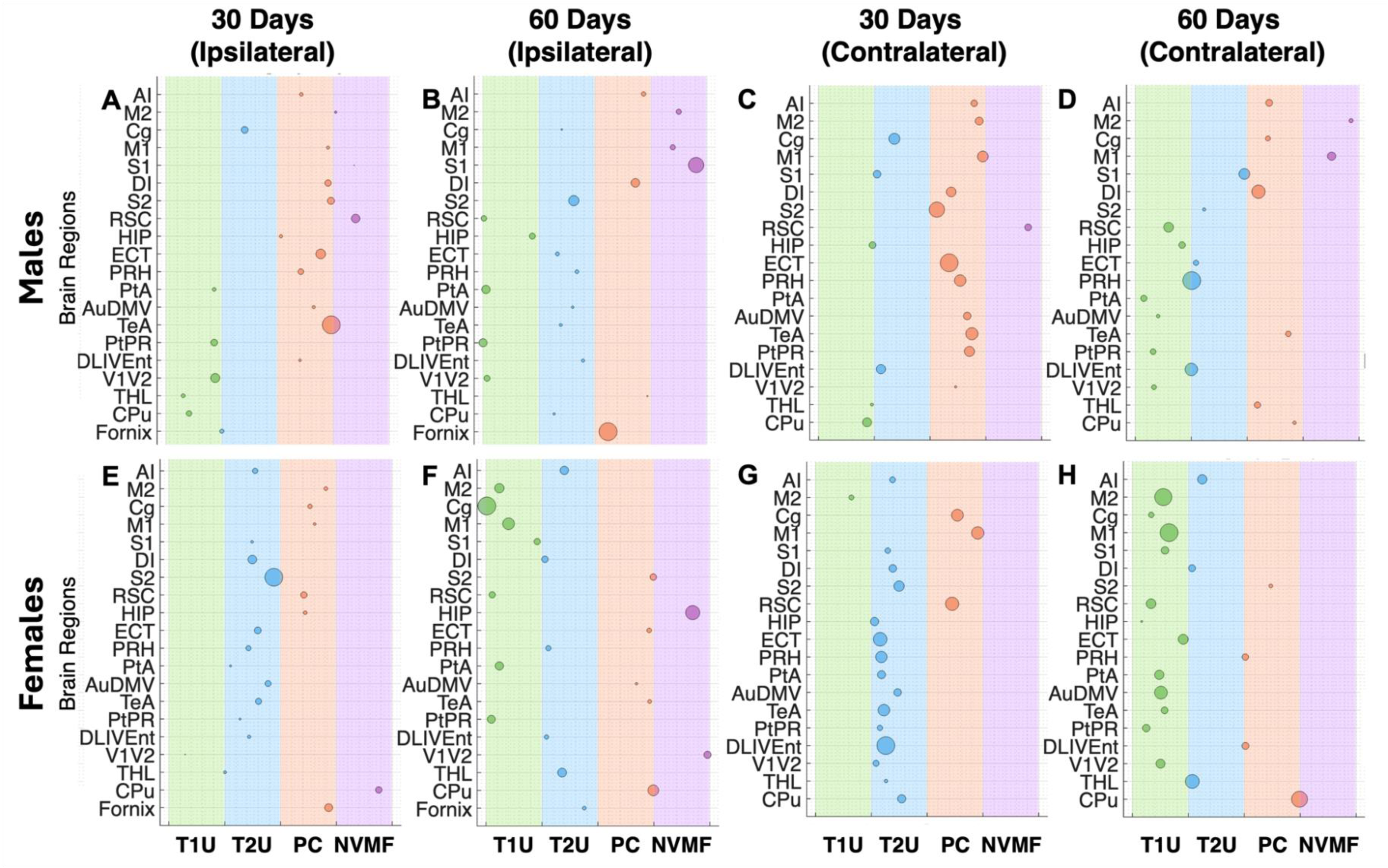
Neuro-Metabolic and Vascular Dysregulation (MVD) Migration Charts of Ipsilateral (ILH) and Contralateral (CLH) Hemispheres. MVD Migration charts (MMCs) generated from the 2D Cartesian MVD plots for males and females for (A, E) ILH at 30-DPI, (B, F) ILH at 60-DPI, (C, G) CLH at 30-DPI, and (D, H) CLH at 60-DPI. MMCs help visualize the trajectories of each brain region (y-axis) across the MVD phenotypes, where circle diameter indicates the magnitude of change (larger circles representing greater deviation of CHI from Sham), and circle position indicates the MVD trajectory location (x-axis). These results indicate that TBI at 30-DPI has distinct effects on males and females across brain regions in both hemispheres, with males exhibiting larger effects than females in regions such as the Temporal Association Cortex (TeA), M2, RSC, and S2. However, by 60-DPI, most of these changes have diminished. For a complete list of brain regions, see Table S1.

### 2.2 Metabolic Connectomics Analysis

#### 2.2.1 Whole-Brain Analysis

Whole-brain MFC analysis was performed at 30- and 60-DPI for both hemispheres and stratified by sex (**Figure S2A/S2B)**. Network organizations were evaluated using whole-brain network metrics. These included degree distribution (number of connections per region), positive and negative node connection strengths (NCS; the sum of the positive or negative weight values for each region, capturing the magnitude of co-activating or anti-correlated relationships), and clustering coefficient (the number of fully connected triangles around a node, based on the weight and sign of the connections, measuring local segregation).^47^

As expected in Sham mice, comparisons between hemispheres in both males (**Figure 4**) and females (**Figure 5**) for the most part do not yield significant differences when comparing the hemisphere across timepoints. These results imply that the brain network integrity of the Sham mice across both time points was maintained. Similarly, in CHI mice for both sexes, few inter-hemisphere comparisons revealed significant changes. However, CHI males at 30-DPI exhibit a significant increase in degree distribution (**Figure 4A**), positive NCS (**Figure 4C**), and negative NCS (**Figure 4D**) in the CLH relative to the ILH. This trend is not observed in females; instead, significant changes occurred across time points. Surprisingly, clustering coefficients showed no significant change in either males (**Figure 4B**) or females (**Figure 5B**). This suggests that the brain reorganizes itself to compensate for changes in local connectivity and metabolic complexity, requiring a more in-depth regional analysis to determine these changes.

**Figure 4.**
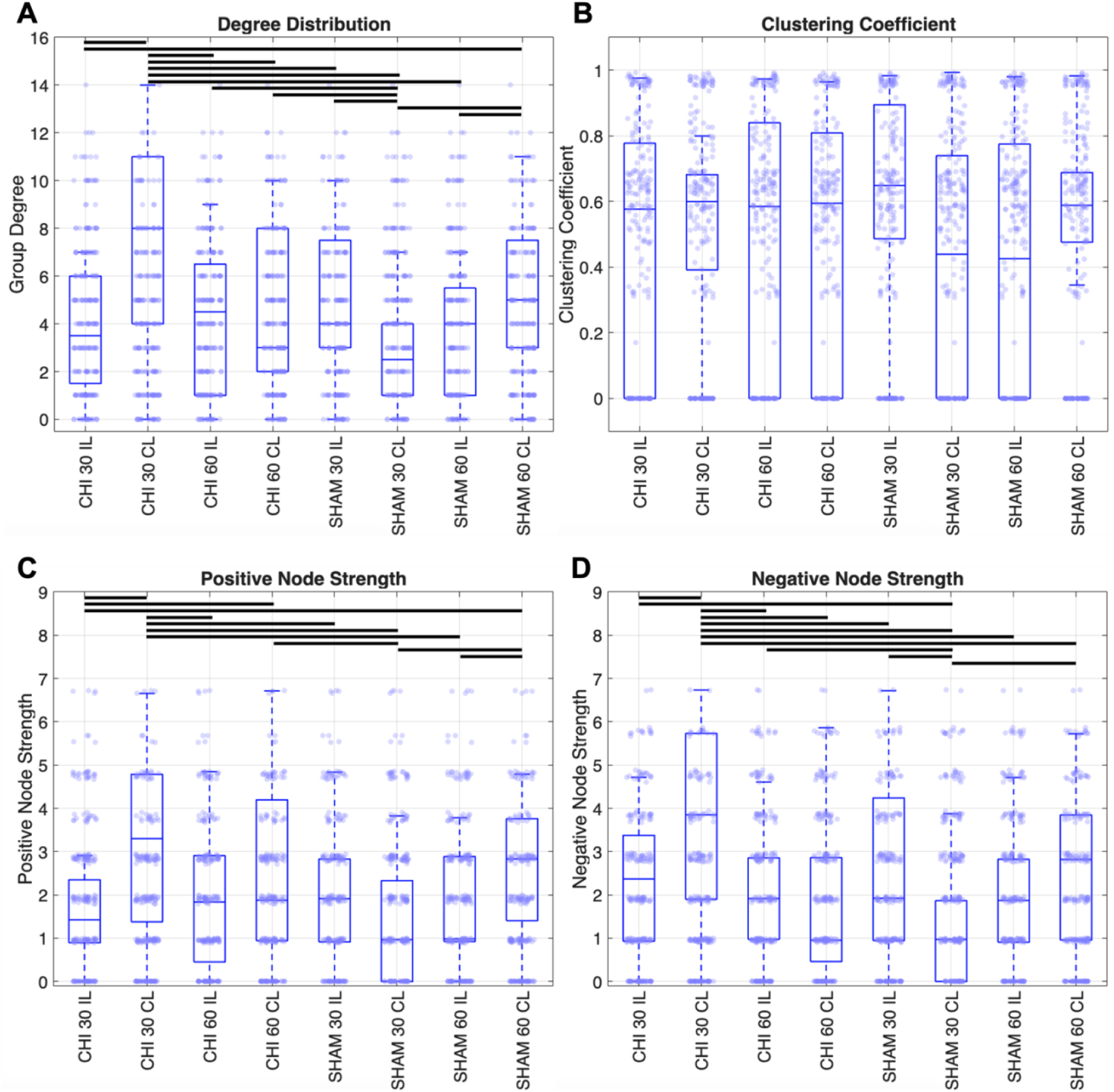
Whole-Brain Network Metabolic Connectomics Analysis for Males. Whole-brain network organization was evaluated using **(A)** degree distribution, **(B)** clustering coefficient, **(C)** positive node connection strengths, and **(D)** negative node connection strengths. Data are shown as Box-Whisker plots with individual values overlaid; black bars indicate p<0.05 via Student’s t-Test. For each metric, the first four Box-Whisker plots correspond to CHI mice, and the last four to Sham mice. Within each group, consecutive pairs represent 30-DPI and 60-DPI, which alternate between ipsilateral (IL) and contralateral (CL). At 30-DPI, CHI mice show decreases in degree distribution, clustering coefficient, and positive node strengths in IL and increases in CL compared to Sham. At 60-DPI, most of these differences are attenuated. These network-level changes are consistent with brain network reorganization after CHI, followed by partial recovery toward Sham-like organization by 60-DPI.

**Figure 5.**
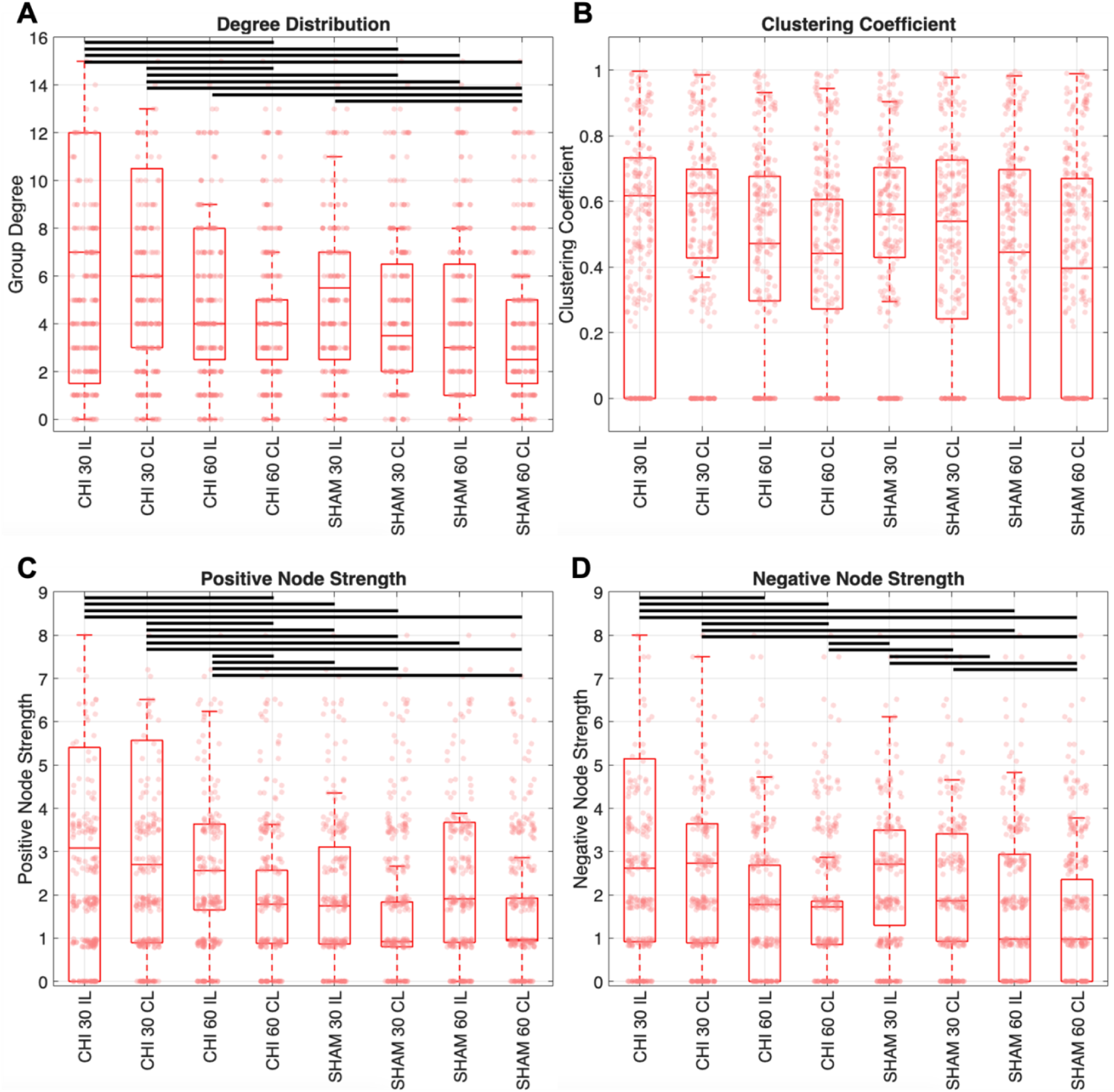
Whole-Brain Network Metabolic Connectomics Analysis for Females. Whole-brain network organization was evaluated using **(A)** degree distribution, **(B)** clustering coefficient, **(C)** positive node connection strengths, and **(D)** negative node connection strengths. Data are shown as Box-Whisker plots with individual values overlaid; black bars indicate p<0.05 via Student’s t-Test. For each metric, the first four Box-Whisker plots correspond to CHI mice, and the last four to Sham mice. Within each group, consecutive pairs represent 30-DPI and 60-DPI, which alternate between ipsilateral (IL) and contralateral (CL). In contrast to male mice, at 30-DPI, female mice exhibit increases in these metrics in both hemispheres at different rates. At 60-DPI, as in male mice, the differences attenuated across metrics. In female mice, these metrics suggest that females exhibit earlier or more stable network reorganization than males.

When comparing changes between CHI and Sham mice, distinct sex-dependent and hemisphere-dependent changes can be observed across both sexes. At 30-DPI, male mice (**Figure 4**) exhibit decreases in degree distribution, clustering coefficient, and positive NCS in CHI ILH, while these metrics increase in the CLH compared to their Sham counterparts. This indicates that ILH exhibited a weaker connectivity profile, characterized by reduced local clustering and greater network homogeneity, whereas CLH displayed an enhanced connectivity profile, accompanied by stronger local segregation and greater nodal organization, which represents a compensatory effect. Conversely, female mice (**Figure 5**) exhibit higher degree distributions, clustering coefficients, and positive NCS in both hemispheres at different rates, consistent with altered metabolic network organization.

At 60-DPI, inter-hemispheric differences within the CHI mice were reduced across most network metrics in both sexes, except for degree distribution in males and positive NCS in females. Compared with their respective Sham counterparts, CHI mice also showed attenuated differences across most network metrics, and this convergence toward Sham was more pronounced in females than in males. Combined, these patterns suggest partial recovery of the brain network toward pre-injury organization by 60-DPI, with females exhibiting earlier or more stable network reorganization than males. However, persistent CHI-Sham disparities on both hemispheres across several metrics indicate that the brain does not fully recover from the injury.

In summary, the findings indicate that network integrity was maintained overtime in Sham mice of both sexes, whereas CHI cases exhibited significant sex-dependent changes in several network metrics in both hemispheres and across both time points.

#### 2.2.2 Regional-Brain Analysis

Regional MFC analysis was performed using metabolic covariance matrices (**Figures S3/S4**) to evaluate network-level changes across the sixteen possible groups, defined by all combinations of sex (females/males), CHI/Sham status, hemisphere (ILH/CLH), and time points (30-/60-DPI). These matrices quantify the extent to which metabolic activity covaries across brain regions, reflecting coordinated inter-regional metabolic organization at the network level; higher covariance values indicate stronger metabolic coupling between regions, while lower values suggest reduced coordination.^47,49^

The analysis was performed by constructing regional MFC modules (module) using the metabolic covariance matrices and multi-resolution consensus clustering (MRCC),^50^ a method for identifying communities across spatial scales and quantitatively determining a consensus partition.^50^ In addition, matrix similarity was calculated using the Frobenius norm (FRO),^51,52^ a matrix norm that can be used to measure the distance between two matrices by applying it to their difference, with smaller values indicating greater similarity and no upper-level bound.^51,52^

Metabolic covariance matrices (**Figures S3/S4**) are symmetric matrices in which each element reflects the pairwise correlation between brain regions, with diagonal elements corresponding to self-correlations. Black squares denote module identified using MRCC. Clinical and preclinical males start with more integrated networks (fewer modules) that degrade under ADRD pathophysiological conditions (more modules), whereas females follow the opposite trend (from fractured to integrated networks).^38,47^ Analyzing these matrices reveals that, regardless of sex, the number of module varies across hemispheres and time points in CHI mice. In Sham mice, females exhibit consistent module counts across hemispheres at matched time points (e.g., ILH and CLH at 30-DPI), while males show hemispheric differences, although the relative pattern is preserved (e.g., CLH has more modules than ILH). Notably, module composition is not stable across conditions, and regions dynamically reassign between module. For instance, in male CHI mice at 30-DPI, M1 clusters with 12 regions (e.g., S1, S2, TeA) in ILH, but clusters with 14 regions (e.g., Entorhinal Cortex (ECT), Perirhinal Cortex (PRH), Caudate Putamen (CPu)) in CLH. Surprisingly, female CHI CLH at 30-DPI showed a single MFC module, suggesting a relatively homogeneous network organization with weak modular segregation. These findings suggest metabolic network reorganization driven by both injury and aging.

The complexity of the results makes simultaneous comparison using the module across hemispheres, time points, and experimental conditions challenging. However, because these matrices are symmetric, they can be reordered to facilitate comparisons by defining one group as the reference and the other as the comparison group, per our previous work.^47,49^ In this study, we combined two matrices such that the upper triangular matrix represents the metabolic covariance matrix of the comparison group, while the lower triangular matrix represents that of the reference group. Subsequently, the module of the reference group is applied. Then, three different comparisons are made: CHI relative to Sham (**Figure 6**), Sham relative to CHI (**Figure S5**), and ILH relative to CLH (**Figure S6**). This provides insight into changes in the interregional relationships associated with injury and aging.

**Figure 6.**
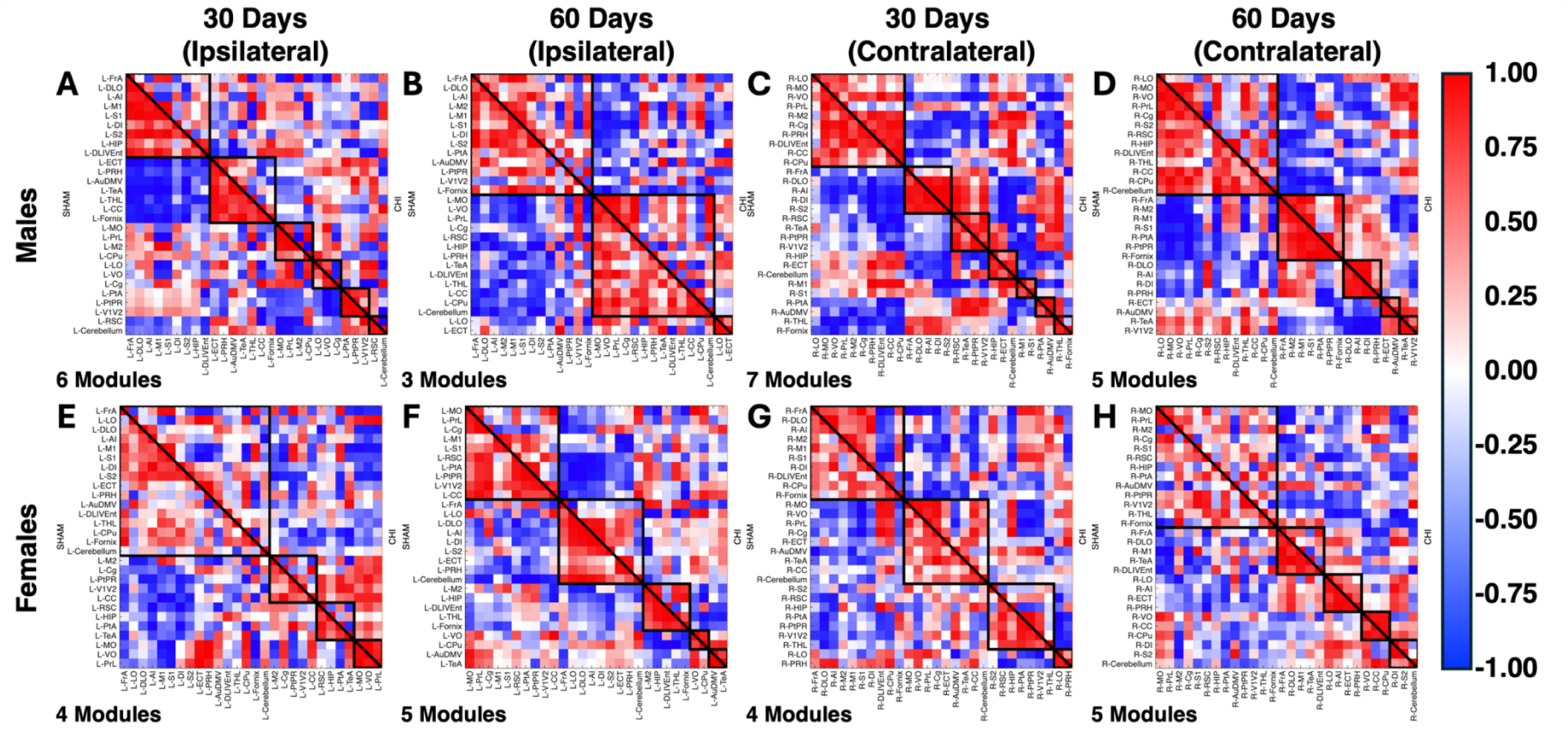
Regional-Brain Analysis of CHI Relative to Sham. Regional metabolic functional connectomics (MFC) analysis using metabolic covariance matrices for **(A-D)** males and **(E-H)** females for **(A, E)** ILH at 30-DPI, **(B, F)** ILH at 60-DPI, **(C, G)** CLH at 30-DPI, and **(D, H)** CLH at 60-DPI. The upper triangular matrix is the comparison group (CHI), and the lower triangular matrix is the reference (Sham). Black outlines denote regional MFC modules (RMCM) identified using multi-resolution consensus clustering (MRCC) for Sham at 30-DPI. Data with warmer colors show positive correlations, while cooler colors show negative correlations. Across both sexes **(A, E)**, CHI ILH at 30-DPI shows more disorganized networks than Sham, indicated by a mix of positive and negative correlations within RMCMs. In this context, positive correlations indicate coupled metabolic activity, while negative correlations suggest inhibited or functionally uncoupled regions. This effect is less apparent in CHI CLH at 30-DPI **(B, F)** and partially reduced by 60-DPI **(B, D, F, H)**. These findings suggest recovery toward a pre-injury state and align with results from MVD and whole-brain analyses.

Analysis of CHI relative to Sham within the same sex, time point, and hemisphere shows that CHI ILH at 30-DPI (**Figures 6A/6E**) exhibits a more disorganized networks (mixture of positive and negative correlations rather than predominantly positive) within the RMCMs than its Sham counterparts. In this context, positive correlations indicate coordinated metabolic excitatory activity, and negative correlations indicate inhibited or functionally uncoordinated regions. A similar but less pronounced effect appears in CLH matrices (**Figures 6C/6G**) in both sexes. At 60-DPI (**Figures 6B/6D/6F/6H**), these differences are partially reduced, suggesting that CHI mice are recovering to a pre-injury state, and are consistent with findings from MVD and whole-brain analyses.

By exchanging the reference and comparison groups (Sham relative to CHI; **Figure S5**), Sham networks appeared less organized at 30-DPI. However, this may be due to the module organization, generated under CHI conditions, which utilizes the metabolic complexity ordering for regions. In addition, these modules also suggest that regions are clustering in a more pathophysiological structure, thus forcing brain regions to cluster together as a compensatory mechanism. Interestingly, as in the previous comparison, brain metabolic networks begin to subtly resemble one another at 60-DPI as measured by FRO.

Comparison across hemispheres (ILH relative to CLH; **Figure S6**) reveals a similar pattern to previous comparisons at both CHI time points. At 30-DPI, ILH shows disorganization within modules, indicating that CHI-induced injury drives these metabolic alterations. By 60-DPI, ILH and CLH show similarity for both sexes and hemispheres, consistent with brain functional recovery. As expected, these alterations remain minimal in Sham mice, as both hemispheres are comparable at both time points.

Matrix similarity quantification across time points (**Table S2A/S2B**) for both sexes and hemispheres using FRO demonstrated that CHI matrices relative to Sham at 30-DPI exhibited lower similarity than their 60-DPI counterparts. FRO values at 30-DPI were higher (males: ILH/CLH=24.22/20.10, females: ILH/CLH=21.00/20.42) than at 60-DPI (males: ILH/CLH=19.03/19.36, females: ILH/CLH=18.66/15.84), indicating greater divergence at the earlier DPI. Fractional percentage changes (FPCs; **Table S2C**) between time points showed a substantial reduction in male ILH (−21.44%) and minimal change in CLH (−3.70%), while females showed large reductions in both hemispheres (ILH=-11.14%, CLH=-22.41%). These findings suggest sex-dependent patterns of network reorganization post-injury, in which males show a unilateral adaptation in ILH relative to CLH, while females show bilateral adaptation. Inverting the reference (Sham relative to CHI) returns the same results, which is expected since the changes for FRO are sign invariant.^51,52^

Comparison between hemispheres within the CHI mice further emphasized sex-dependent differences. Males exhibited a substantial decrease over time (FPC=-46.33%/FRO 30-DPI=23.93/FRO 60-DPI=12.84), while females showed a smaller reduction (FPC=-9.31%/FRO 30-DPI=14.99/FRO 60-DPI=13.59). Consistent with previous reports,^26–28^ these findings support the existence of sex-dependent mechanisms underlying metabolic recovery. In contrast, Sham mice exhibited small divergence across time (males: FPC=-11.83%/FRO 30-DPI=18.45/FRO 60-DPI=16.35; females: FPC=1.35%/FRO 30-DPI=16.18/FRO 60-DPI=16.40) compared to the changes observed in CHI mice, indicating preserved interhemispheric similarity in the absence of injury.

In summary, these results show that, regardless of the reference modules, this approach can detect temporal and hemispheric alterations in the brain networks, although the directionality of change (i.e. FPC), depends on the selected reference. At the regional level, these findings indicate that the metabolic brain networks are reorganizing to compensate for, and recover connectivity towards, their pre-injury state, with females exhibiting earlier stabilization or partial recovery than males. However, these differences begin to diminish at 60-DPI. Collectively, whole-brain and regional analyses demonstrate a temporal, sex- and injury-location-dependent evolution in metabolic complexity.

### 2.3 Region-Set Enrichment Analysis

Region-set enrichment analysis (RSEA) was conducted by clustering brain regions using the modules derived from male and female Sham mice at 30-DPI for each hemisphere via MRCC, per our previous methods (**Figure S2C)**.^47,49^ For all RSEA heatmaps, negative values represent a fractional percentage reduction in metabolic demand for a given functional category in the comparison group relative to the reference, while positive values indicate a relative increase, and values closer to 0 represent that both groups are similar to each other.

At 30-DPI, male CHI mice exhibited predominantly negative changes in the ILH, with most modules in the learning, motor, perception, and sensory categories showing reduced metabolic demand relative to Sham (**Figure 7A**). In contrast, the CLH in males displayed a mixture of positive and negative changes across the same categories, indicating greater heterogeneity during reorganization (**Figure 7C**). In females, both hemispheres demonstrated a mixture of increases in the learning, integration, and perception categories, while motor and sensory categories were consistently reduced across modules (**Figures 7E/7G**). Notably, module #1 in females showed a marked increase in the learning category, accompanied by concurrent changes in motor, perception, and integration, suggesting a learning-related network reorganization alongside suppression of sensorimotor networks at 30-DPI. These sex-specific patterns suggest that males primarily exhibit broad downregulation of metabolic demand, while females appear to engage in compensatory upregulation.

**Figure 7.**
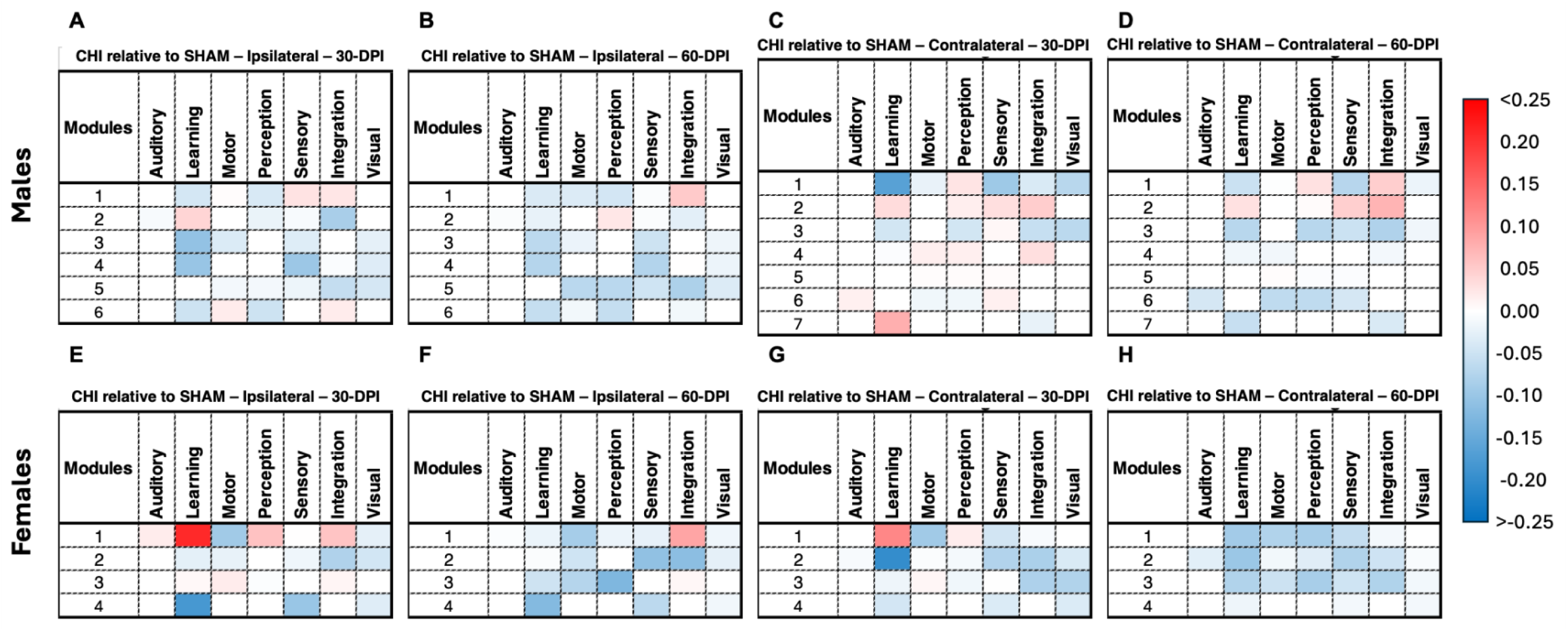
Region-Set Enrichment Analysis of CHI relative to Sham using 30-DPI Sham RMCM. Region-set enrichment analysis (RSEA) heatmaps are shown for **(A-D)** males and **(E-H)** females. These are split as follows: **(A, E)** ILH at 30-DPI, **(B, F)** ILH at 60-DPI, **(C, G)** CLH at 30-DPI, and **(D, H)** CLH at 60-DPI. RMCM were obtained using MRCC on the Sham mice at 30-DPI for each hemisphere. These heatmaps show metabolic changes across seven functional categories, based on ^18^F-FDG readouts. Data with warmer colors shows positive fractional percentage changes, while cooler colors show negative fractional percentage changes. In these RSEA heatmaps, positive changes indicate a fractional percentage increase in metabolic demand for a given functional category in the comparison group (CHI) relative to the reference (Sham), while negative changes indicate a relative decrease, and values near 0 indicate similarity between groups. At 30-DPI, males exhibit predominantly negative changes in the ILH in learning, perception, and sensory categories. Females exhibit a mixture of increases and decreases in metabolic changes across these categories, suggesting compensatory upregulation. By 60-DPI, RSEA changes were diminished in both sexes, indicating partial functional recovery or stabilization of metabolic complexity. However, CHI-induced injury persists, with subtle alterations observed across both time points and hemispheres.

At 60-DPI, RSEA alterations were attenuated in both sexes, with most modules in ILH modules in male (**Figure 7B**) and female (**Figure 7F**) CHI mice showing reduced deviation (lighter blue/red relative to 30-DPI) from Sham, consistent with partial functional recovery or stabilization of metabolic demand. CLH displayed a similar pattern across module and functional categories (**Figures 7D/7H**). Therefore, the subtle changes indicate that that 60-DPI CHI-induced injury persists at low levels, even when there is some evidence of recovery in the brain network and metabolic complexity. Across both time points and hemispheres, auditory and visual categories showed only subtle alterations, which may reflect either the relative resilience of these functional systems to CHI injuries or their distance from the injury location.

When comparisons are made between Sham relative to CHI using CHI 30-DPI module (**Figure S7**), both sexes exhibit large increases across most functional categories, except for the auditory category, which shows subtle changes. Within-hemisphere comparisons from 30- to 60-DPI also reveal increases in most function categories, but the extent of these changes varies by module. These findings indicate that the metabolic demand in CHI mice is reduced (hypometabolic) compared to Sham mice at 30-DPI, suggesting that CHI mice may remain in a partial recovery phase with reduced metabolic activity, reinforcing the idea that CHI-induced changes dampen overall network-level metabolic demand. At 60-DPI, some functional categories show evidence of partial recovery (e.g., modules 4 and 5 in females; **Figure S7F**). However, comparisons using the CLH CHI modules are difficult to perform due to the reduced number of modules.

When comparisons are made between hemispheres using the CLH modules at 30-DPI (**Figure S8**), results show CHI- and Sham-dependent changes. In CHI mice (males: **Figure S8A-B**; females: **Figures S8E-F**), ILH shows metabolic decreases across multiple functional categories and modules in both sexes. In contrast, Sham mice (males: **Figure S8C-D**; females: **Figures S8G-H**) show more subtle differences. The patterns at 30- and 60-DPI are similar, suggesting that CHI induces a persistent reduction in metabolic demand in ILH relative to CLH, that does not recover by 60-DPI.

Collectively, these RSEA results highlight sex-, hemisphere-, and time-dependent variations in functional network organization following CHI. Male mice exhibit a pronounced early reduction in learning, perception, and sensory categories in the ILH at 30-DPI, which largely normalizes by 60-DPI relative to Sham. In contrast, females showed early hemisphere-specific increases in learning, perception, and integration categories, superimposed on reductions in sensorimotor categories, which also return toward Sham-like patterns by 60-DPI. Overall, CHI induces lateralized network restructuring, with more pronounced widespread downregulation in males, while females exhibit both metabolic upregulation and downregulation.

## 3. DISCUSSION

This study evaluates the construct and face validity of our CHI model in middle-aged C57BL6/J mice and leverages our novel MVD and MFC analysis methods to assess regional phenotypes (**Figure 2**) in key brain regions on the ILH (i.e. S2, RSC, DLIVEnt, M1, M2, Cg, TeA). These regions showed accelerated MVD trajectories, as indicated by the size and location of the circles along the x-axis in the MMC (**Figure 3**). As expected, the ILH exhibits greater MVD compared to the CLH. Analysis of the MMC reveals sex-specific differences at 30-DPI, where males (**Figure 3A-D**) showed a greater number of regions in the PC phenotype, indicating severe dysfunction, while females (**Figure 3E-H**) had greater number regions in the T2U phenotype, and differences are present across both brain hemispheres. By 60-DPI, most regions in both sexes and brain hemispheres transition to the T1U phenotype, suggesting a capacity for MVD recovery toward a pre-injury state. The normal MVD sequence is T1U◊T2U◊PC◊NMVF, and has been shown to be initiated by acute neuroinflammation, that progresses to a chronic state, yielding the PC phenotype.^36,48^ Overall, while sex influences the rate and pattern of recovery following TBI, these findings indicate that, regardless of sex and at the time of assessment, the brain demonstrates a substantial ability to partially recover MVD function over time. These findings are consistent with current research, which shows that TBI induces pronounced, region-specific MVD dysfunction in the hemisphere closest to the injury site, with a characteristic pattern of initial dysregulation followed by recovery.^31,53,54^ From the sex-specific response perspective, recent studies report that males experience more severe early dysfunction^55,56^, while females exhibited a distinct recovery profile, potentially linked to hormone-mediated neuroprotection and differential inflammatory responses.^55–58^ Given this, MVD may serve as both biomarker of TBI pathology and recovery, thus supporting its relevance for outcome assessments and individualized therapeutic strategies in both animal models and human patients.^31,53^

From the MFC perspective, our analysis revealed both sex- and hemisphere-dependent changes in network connectivity at whole-brain (**Figures 4/5**) and regional (**Figures S3/S4/6**) levels. Whole-brain analysis (**Figures 4/5**) supports these findings by revealing CHI changes, relative to sham, align with the network alterations observed in migration uncoupling charts (**Figure 3**) and in metabolic covariance matrices (**Figure S3/S4**). At a regional-level (**Figures 6/S5/S6**), male CHI animals generally exhibited more fractured networks at 30-DPI, as indicated by changes in the number of MRCC modules. By 60-DPI, these alterations begin to diminish as measured by FRO (**Table S2**), indicating a network recovery between time points. In contrast, females showed more integrated networks at 30-DPI, which became more fractured by 60-DPI. These results suggest that, following TBI, brain regions reorganize their network structures adaptively to mitigate injury effects and are consistent with patterns observed in mice and humans.^40–42,44,45^

Taken together, MVD and MFC observations support the hypothesis that the brain dynamically reorganizes its metabolic network architecture to compensate for cerebral perfusion and metabolic disruption caused by TBI, with patterns that differ by sex and injury location. Current TBI studies consistently demonstrate disruptions in brain network structure and function, particularly in the ILH regions,^39,59–61^ as showed by alterations in clustering coefficients after injury, with recovery manifesting as normalization of these network metrics over time.^61^ Importantly, these network alterations exhibit both sex- and hemisphere-dependence, supporting the growing consensus that post-TBI recovery mechanisms are influenced by biological sex and intervention strategies.^59–61^

Comparison of RSEA results indicates that our framework detected changes across multiple functional categories following TBI, when compared with Sham mice (**Figure 7**). Males exhibit widespread metabolic downregulation across functional categories on the injury side. By contrast, females show a mixture of both up- and down-regulated metabolic demands across categories. In this context, negative changes reflect a reduction in function, while positive changes indicate an increased metabolic energy demand to sustain functions. Temporal analysis shows that alterations observed at 30-DPI shift towards the Sham counterparts by 60-DPI, suggesting progressive brain recovery. These metabolic changes are likely associated with neuroinflammatory processes following TBI, which have been previously reported.^12,13,16,27^ Our study supports these findings whereby TBI produces both sex- and region-specific changes in brain metabolism, with males exhibiting greater metabolic downregulation and heightened vulnerability to TBI-induced dysfunction compared to females.^27,29,62–64^ Furthermore, the more pronounced early-stage energy dysfunction and downregulation in males is attributed to the more aggressive neuroinflammatory responses and increased neuronal death.^27^ In contrast, in some experimental TBI models, females exhibit differential cerebrovascular responses^26^ and reduced neuroinflammation^65^ compared with males, an effect often attributed to hormonally mediated neuroprotection and differences in inflammatory signaling.^26,65^ These findings, combined with current research,^27,66^ suggest that males may be more susceptible to TBI-induced vascular and metabolic dysfunction than females at early time points, implying that females could possess a greater adaptive response to neuroinflammation. Analyses using different reference groups show similar patterns; Sham relative to CHI (**Figure S7**) shows that CHI-induced changes reduce overall network-level metabolic demand at early stages with partial recovery over time, while ILH relative to CLH (**Figure S8**) reveals a persistent reduction in the metabolic demand over the injury location. These data suggest that age, sex, genetics, and time of injury may be important factors in recovery and residual disease post-TBI and may impact future ND progression.

There is emerging clinical and neuropathological overlap between TBI and ADRD, suggesting that TBI may accelerate individuals into AD-like pathology.^4–7^ In ADRD, our understanding of disease mechanisms has advanced through the use of transgenic^37,67–69^ and knock-in^70^ mouse models that can reflect human phenotypes, including Aβ and tau deposition (key pathological hallmarks of AD),^71^ chronic neuroinflammation,^67^ increased oxidative stress,^68^ and immune,^72^ metabolic,^73,74^ and vascular dysfunctions.^37,69^ However, the 2025 ADRD Summit^8^ highlighted a substantial gap in our understanding of the mechanisms by which TBI may act as a potential driver in the onset and progression of ADRD through gene-by-environment interactions. The MVD alterations observed in our study parallel those reported in AD mouse models bearing *Apolipoprotein E*4 risk alleles, although these effects are predominantly observed in females.^36^ Moreover, the MFC patterns identified in CHI mice are consistent with prior clinical and preclinical findings in ADRD reported by our lab and others,^38,47,49^ in which male controls exhibit more integrated networks that fracture with disease onset and progression, while females begin with more fractured networks that become more integrated under pathological conditions.^38,47,49^ Collectively, these converging findings highlight the need for strong collaboration between TBI and ADRD researchers to bridge the gap in understanding post-TBI pathology and its influence on onset and progression of ND.

As with any research, it is important to acknowledge the limitations of this study. First, the study focused on the impact of a CHI delivered to a single hemisphere (left) and did not test whether injuries to the opposite hemisphere, different cortical locations, or repeated injuries would produce distinguishable effects. Second, behavioral testing was not performed, and all imaging data were acquired under resting conditions; consequently, there is limited insight into how these neuroimaging findings relate to behavioral outcomes or how the brain responds under conditions that require increased metabolic activity or vascular flow. Finally, because perfusion and metabolic images were acquired on different days, simultaneous assessment of both readouts was not possible, precluding a fully integrated evaluation of the temporal relationship between blood flow and metabolism.

In summary, this study demonstrates that the data processing pipeline is a reproducible framework that produces a reliable and quantifiable CHI phenotype, establishing a foundation for diagnostic and monitoring in TBI research. The findings further suggest sex-dependent alterations in neurometabolic and vascular regulation, as well as in brain metabolic connectivity networks, following CHI, resulting in temporal changes in brain networks and functional alterations. Consequently, this framework provides a basis for evaluating the impact and interaction of TBI on other pathological conditions (e.g., ADRD, Parkinson’s disease) in both temporal and long-term perspectives.

## 4. METHODS

### 4.1 Dataset

All studies were conducted in accordance with the approval of the Institutional Animal Care and Use Committee (IACUC) of the University of California Irvine Medical School and the National Research Guide for the Care and Use of Laboratory Animals, and adhered to the ARRIVE guidelines (A. Obenaus moved to University of California Riverside after these data were collected).^75^ Male and female C57BL/6J (B6) mice at 8-months-old were randomized by sex on each injury day. Mice were administered a single TBI over the left mid-brain using the closed-head injury (CHI) protocol.^76,77^ Sham mice underwent the same procedure without the impact (N=4 CHI males, N=5 CHI females, N=4 sham males, N=5 sham females). Mice were then followed for 30- and 60-days post-injury (DPI) for metabolism and perfusion.

Brain perfusion and cerebral blood flow (CBF) were captured using dynamic contrast enhanced MRI (DCE-MRI) images were acquired as a temporal series of a T1-weighted gradient echo flow compensated (GEFC) images at 3.97 sec/image with the following image parameters: FA (15); TE (3.5ms); TR (61.7ms); 100 repetitions; 4 slices at the level of the dorsal hippocampus, 1mm slice thickness. A pre-contrast baseline of 1.5min was acquired and then mice were intravenously injected with Gadolinium (Dotarem, 0.1mmol/kg) and images were acquired for an additional 3.5min. CBF maps were computed using Jim software (XINAPSE, 9.0) where arterial input voxels automatically derived. All arterial input curves were assessed manually and then CBF maps were computed using a two-compartment model.

Brain metabolism was evaluated using ^18^F-FDG PET. All PET/CT imaging was performed on an Inveon PET/CT scanner (Siemens Medical Systems, Knoxville, TN). Awake, overnight-fasted (18 hours) mice (maintained in an enclosure at 25-27°C) were administered 3.9-4.6 MBq of ^18^F-FDG intraperitoneally. Following injection, mice were returned to their cages at 25-27 °C with free access to water. Forty minutes post-injection, mice were anesthetized with 3% isoflurane, placed in the supine position in a mouse holder, and maintained with 3% isoflurane in the PET/CT scanner. Sequential CT and PET imaging of the mice were carried out in the same bed position. A 7.5-minute-long CT scan for attenuation correction and anatomical delineation of PET images was followed by a 30 min-long static PET scan. Upon completion of imaging, mice were returned to their home cages for recovery. PET data were reconstructed as 128×128×159 matrices with a trans-axial pixel of 0.776 mm and a slice thickness of 0.796 mm using the OSEM3D algorithm (two OSEM iterations, 18 MAP iterations). PET images were corrected for random coincidences, attenuation, and scatter according to the manufacturer’s instructions. All images were calibrated in units of Bq/cm3 by scanning a Ge-68 cylinder (6 cm diameter) with known activity. CT images were spatially transformed to align with the reconstructed PET images. CT projections were acquired with the detector-source assembly rotating over 360°, collecting 180 projections at 80 kV, 500 mA, and 23.46 μm pixel resolution. CT images were reconstructed using a Feldkamp reconstruction protocol and a Shepp-Logan filter with a cutoff at the Nyquist frequency and a binning factor of 2, resulting in an image matrix of 512×512×1008 and a voxel size of 0.052 mm^3^.

### 4.2 Study Workflow

The study workflow diagram (**Figure 1B**) outlines the experimental timelines, procedures, sexes, and readouts used. Briefly, 8-month-old male and female mice (n=4-5/sex/treatment) received a single CHI TBI (sTBI) over the left M1 and M2 cortex, MRI and PET imaging were conducted at 30- and 60-DPI. MVD, metabolic connectomics, and RSEA analyses methods were conducted on datasets from both time points.

### 4.3 Neurometabolic and Vascular Dysregulation Analysis Framework

As described in our previous work,^36^ imaging data (DCE-MRI, ^18^F-FDG PET) were co-registered to the Paxinos-Franklin atlas,^78^ which contains 40 bilateral regions over the region covered by the DCE-MRI images. MVD analyses were performed by extracting mean cerebral blood flow for each bilateral brain region from the registered DCE-MRI as a measure of regional perfusion and the registered ^18^F-FDG PET as a surrogate of cerebral metabolism.^79^ To minimize the dose effect, scanner variability, and brain uptake normalization, regional intensities values were ratioed to the corpus callosum.^80^ Similarly, perfusion regional values were also ratioed to the corpus callosum to maintain the same reference region across both modalities. Subsequently, 30- and 60-DPI CHI values were converted to z-scores relative to Sham 30- and 60-DPI, respectively. Regional z-scored data were projected onto a 2D Cartesian plane, where the x-axis represents the z-scored perfusion and the y-axis representing the z-scored metabolism, as measured by their respective surrogate readouts. This mapping yielded four possible MVD phenotypes: Type 1 Uncoupling (↑ perfusion, ↓ metabolism; T1U), Type 2 Uncoupling (↓ perfusion, ↑ metabolism; T2U), Prodromal Coupled (↑ perfusion, ↑ metabolism; PC), and Neuro-Metabolic-Vascular Failure (↓ perfusion, ↓ metabolism; NMVF). Lastly, for improved MVD monitoring the Cartesian space regional data were transformed into MMC as described in our previous clinical work.^48^ Graphical depiction of this process is shown in **Figure S1**.

### 4.4 Metabolic Functional Connectomics Analysis

Metabolic functional connectomics (MFC) analysis was performed using the SUVR ^18^F-FDG data, in which bilateral brain regions (56 total, 28 for each side) were ratioed relative to the whole brain, following established procedures.^81^ SUVR values were analyzed using open-source packages including Brain Connectivity Toolbox (BCT), generalized Louvain modularity, hierarchical consensus clustering, and CovNet.^49^ As previously described,^47,49^ each of the eight possible groups (all possible combinations between CHI/Sham, ILH/CLH, and 30-/60-DPI) produced a covariance matrix of size 28-by-28 representing the similarity between regions in the ^18^F-FDG readouts across all region pairs. In addition, the following network measures were computed for each covariance matrix: (1) network density, indicating the number of connections; (2) nodes degree, representing the number of connections per region; (3) positive and negative nodal strengths, defined as the sum of positive or negative covariance values for each region; and (4) clustering coefficient, quantifying the number of fully connected triangles around a node based on connection weight and sign. Modularity maximization was performed using multi-resolution consensus clustering (MRCC) to identify groups of nodes in a network from an intra-group connectivity perspective, as described in our previous work.^47,49^ These modules were subsequently used to assess alterations in module connectivity within each group. Graphical depiction of this process is shown in **Figures S2A/S2B**.

### 4.5 Regional-Set Enrichment Analysis

Region-set enrichment analysis (RSEA) is a technique developed in our lab to quantitatively detect brain functional changes by analyzing regional metabolic readout variations between experimental and reference groups.^47^ This method works by weighting changes in each region (experimental group minus the reference group) and then associating these changes to functional-set enrichment categories (FSEC), which include: auditory, learning, motor, perception, sensory, integration, and visual domains. Using the modules identified by MRCC in the metabolic connectomics analysis, modular FSEC scores were generated for each experimental group relative to the reference group by summing the columns corresponding to the regions within each module. Graphical depiction of this process is shown in **Figure S2C**.

### 4.6 Statistical Analysis

Statistical analysis of at-risk regions was evaluated using Student’s t-test (p<0.05), without Bonferroni correction since regional independence was maintained (i.e. region “x” vs. region “x” between experimental stage and the reference stage),^82,83^ where MVD phenotype was determined by the Cartesian location of glycolytic metabolism and perfusion z-scores per our previous works.^36,48^ Significant regions were then projected onto Allen MRI Temple object maps to provide spatial localization, where the sign of the z-score was applied to the p-value to indicate directionality of change (i.e. 0<z increasing, z<0 decreasing). Whole-brain connectomics metrics significance testing between time points was determined via Student’s t-test with a false discovery rate (FDR) of 0.05. Analysis was performed in MATLAB R2024a.

## ACKNOWLEDGEMENTS

The authors wish to thank NIH (NIA and NINDS) for their generous support of this work (RF1NS1380 and T32AG071444).

## AUTHOR’S CONTRIBUTIONS

JAKCC performed MVD, metabolic functional connectomics, and region-set enrichment analyses, and wrote the paper. SAP performed image processing, registration, and analysis on the images. AJ performed image processing of the MRI images and wrote sections of the paper. JY and MB performed the live phase for the TBI and imaging studies. AO and PRT designed the experiments, performed MVD and connectomics analysis, and wrote sections of the manuscript. All authors reviewed and approved the manuscript prior to publication.

## DATA SHARING

The datasets used in this study will be made available upon reasonable request to the corresponding author.

## ETHICS COMMITTEE APPROVAL

All studies were carried out in accordance with institutional and federal guidelines for Animal Care and Use. The Animal Use Protocol #23-067 (formerly #20-110) was approved by the Institutional Animal Care and Use Committee of the University of California, Irvine.

