## Supplemental Materials for "Neuro-Metabolic and Vascular Dysregulation are Functional Biomarkers of Acquired Brain Injury"

### Tables of Contents

**Supplemental Table S1 – Brain Regions**

| Abbreviation | Region Name |
|---|---|
| AI | Agranular Insular Cortex (Doral-Ventral) |
| AuDMV | Auditory Cortex (Dorsal, Medial, Ventral) |
| CC | Corpus Callosum |
| Cg | Cingulate Cortex |
| Cpu | Caudate Putamen (Dorsal Striatum) |
| DI | Dysgranular Insular Cortex |
| DLIVEnt | Dorso-Lateral Intermediate Entorhinal Cortex |
| DLO | Dorsolateral Orbital Cortex |
| ECT | Entorhinal Cortex |
| Fornix | Fornix |
| FrA | Frontal Association Cortex |
| HIP | Hippocampus (CA1-CA3) |
| LO | Lateral Orbital Cortex |
| M1 | Primary Motor Cortex |
| M2 | Secondary Motor Cortex |
| MO | Medial Orbital Cortex |
| PRH | Perirhinal Cortex |
| PrL | Prelimbic Cortex |
| PtA | Perietal Association Cortex (Lateral-Medial |
| PtPR | Parietal Cortex (Post-Rostral) |
| RSC | Retrospleinal Dysgranular Cortex |
| S1 | Primary Somatosenory Cortex |
| S2 | Secondary Somatosenory Cortex |
| TeA | Temporal Association Cortex |
| THL | Thalamus |
| V1V2 | Visual Cortex (Primary and Secondary) |
| VO | Ventral Orbital Cortex |

**Supplemental Table S2 – Frobenius Norm and Fractional Percentage Changes.**

**A Males — Frobenius Norm**

| Comparison | | | | Reference: Ipsilateral CHI 30-DPI | Ipsilateral CHI 60-DPI | Ipsilateral Sham 30-DPI | Ipsilateral Sham 60-DPI | Contralateral CHI 30-DPI | Contralateral CHI 60-DPI | Contralateral Sham 30-DPI | Contralateral Sham 60-DPI |
|---|---|---|---|---|---|---|---|---|---|---|---|
| Comparison | Ipsilateral | CHI | 30-DPI | 0.00 | 22.63 | 24.22 | 23.85 | 23.93 | 22.80 | 23.92 | 24.08 |
| | | | 60-DPI | 22.63 | 0.00 | 22.29 | 19.03 | 24.55 | 12.84 | 20.89 | 20.33 |
| | | Sham | 30-DPI | 24.22 | 22.29 | 0.00 | 21.96 | 23.98 | 21.18 | 18.45 | 23.76 |
| | | | 60-DPI | 23.85 | 19.03 | 21.96 | 0.00 | 24.39 | 18.01 | 17.90 | 16.35 |
| | Contralateral | CHI | 30-DPI | 23.93 | 24.55 | 23.98 | 24.39 | 0.00 | 25.04 | 20.10 | 26.13 |
| | | | 60-DPI | 22.80 | 12.84 | 21.18 | 18.01 | 25.04 | 0.00 | 19.15 | 19.36 |
| | | Sham | 30-DPI | 23.92 | 20.89 | 18.45 | 17.90 | 20.10 | 19.15 | 0.00 | 20.89 |
| | | | 60-DPI | 24.08 | 20.33 | 23.76 | 16.35 | 26.13 | 19.36 | 20.89 | 0.00 |

**B Females — Frobenius Norm**

| Comparison | | | | Reference: Ipsilateral CHI 30-DPI | Ipsilateral CHI 60-DPI | Ipsilateral Sham 30-DPI | Ipsilateral Sham 60-DPI | Contralateral CHI 30-DPI | Contralateral CHI 60-DPI | Contralateral Sham 30-DPI | Contralateral Sham 60-DPI |
|---|---|---|---|---|---|---|---|---|---|---|---|
| Comparison | Ipsilateral | CHI | 30-DPI | 0.00 | 15.53 | 21.00 | 19.98 | 14.99 | 17.81 | 19.13 | 19.02 |
| | | | 60-DPI | 15.53 | 0.00 | 19.78 | 18.66 | 18.51 | 13.59 | 18.46 | 17.98 |
| | | Sham | 30-DPI | 21.00 | 19.78 | 0.00 | 16.58 | 21.14 | 15.03 | 16.18 | 19.63 |
| | | | 60-DPI | 19.98 | 18.66 | 16.58 | 0.00 | 22.48 | 15.52 | 16.98 | 16.40 |
| | Contralateral | CHI | 30-DPI | 14.99 | 18.51 | 21.14 | 22.48 | 0.00 | 19.35 | 20.42 | 20.85 |
| | | | 60-DPI | 17.81 | 13.59 | 15.03 | 15.52 | 19.35 | 0.00 | 17.51 | 15.84 |
| | | Sham | 30-DPI | 19.13 | 18.46 | 16.18 | 16.98 | 20.42 | 17.51 | 0.00 | 16.69 |
| | | | 60-DPI | 19.02 | 17.98 | 19.63 | 16.40 | 20.85 | 15.84 | 16.69 | 0.00 |

**C Fractional Percentage Changes (PFC)**

| Between Time Points | | Males | Females |
|---|---|---|---|
| CHI relative to Sham | ILH | -21.44% | -11.14% |
| | CLH | -3.70% | -22.41% |
| Sham relative to CHI | ILH | -21.44% | -11.14% |
| | CLH | -3.70% | -22.41% |
| ILH relative to CLH | CHI | -46.33% | -9.31% |
| | Sham | -11.38% | 1.35% |

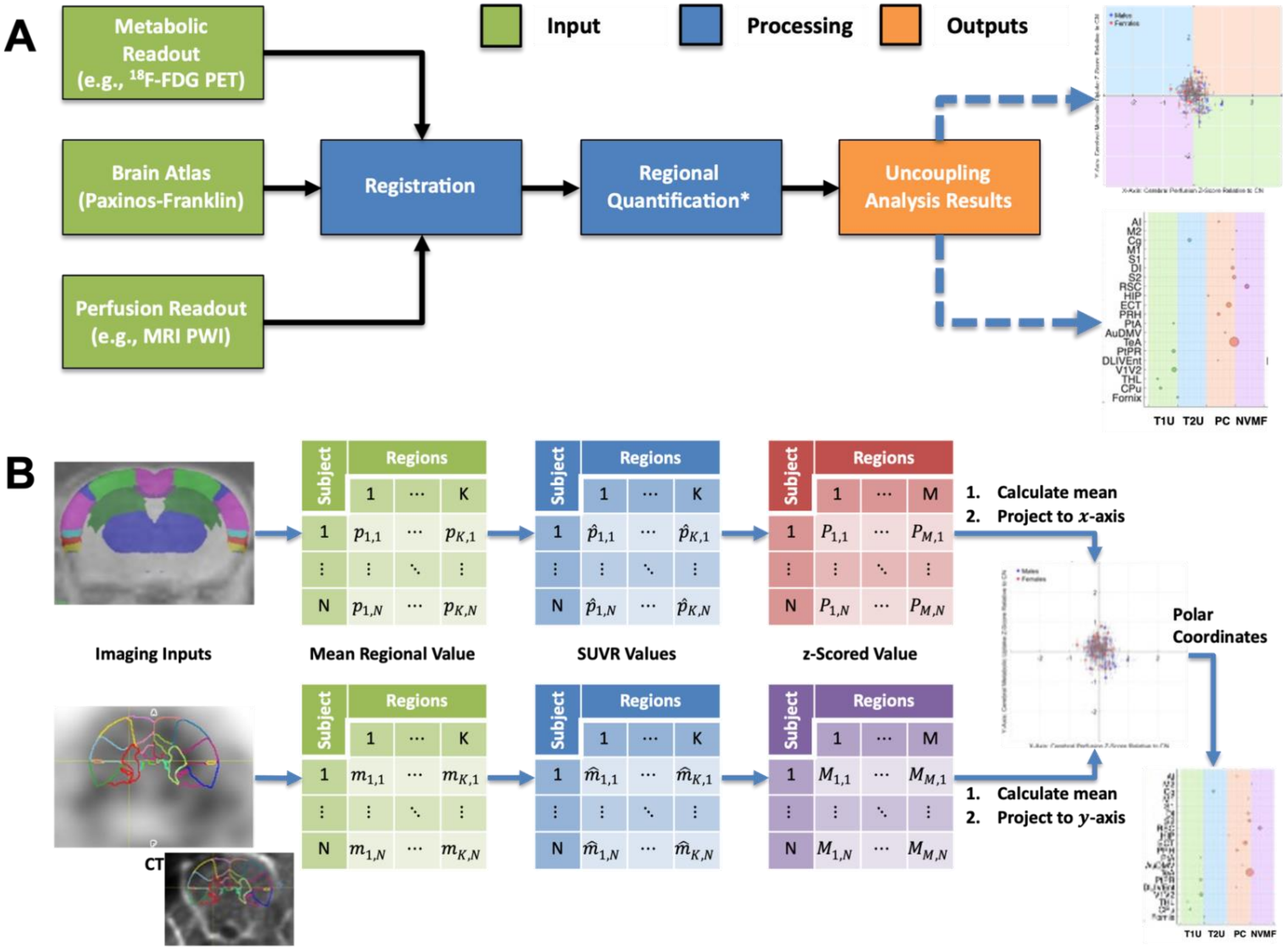

**Supplemental Figure S1 – Analysis Workflow for Neuro-Metabolic and Vascular Dysregulation.**
**(A)** Schematic representation of the neuro-metabolic and vascular dysregulation (MVD) analysis workflow, illustrating how metabolic and perfusion imaging inputs are registered to a brain atlas, regionally quantified, and used to generate 2D MVD plots and MVD Migration Charts. **(B)** Detailed view of data flow through the analysis pipeline, showing extraction of regional values, normalization, and computation of MVD phenotypes and trajectories across the MVD continuum. The MVD analysis workflow quantifies regional metabolic and perfusion changes, determines MVD phenotypes, and tracks the regional trajectories across the MVD continuum.

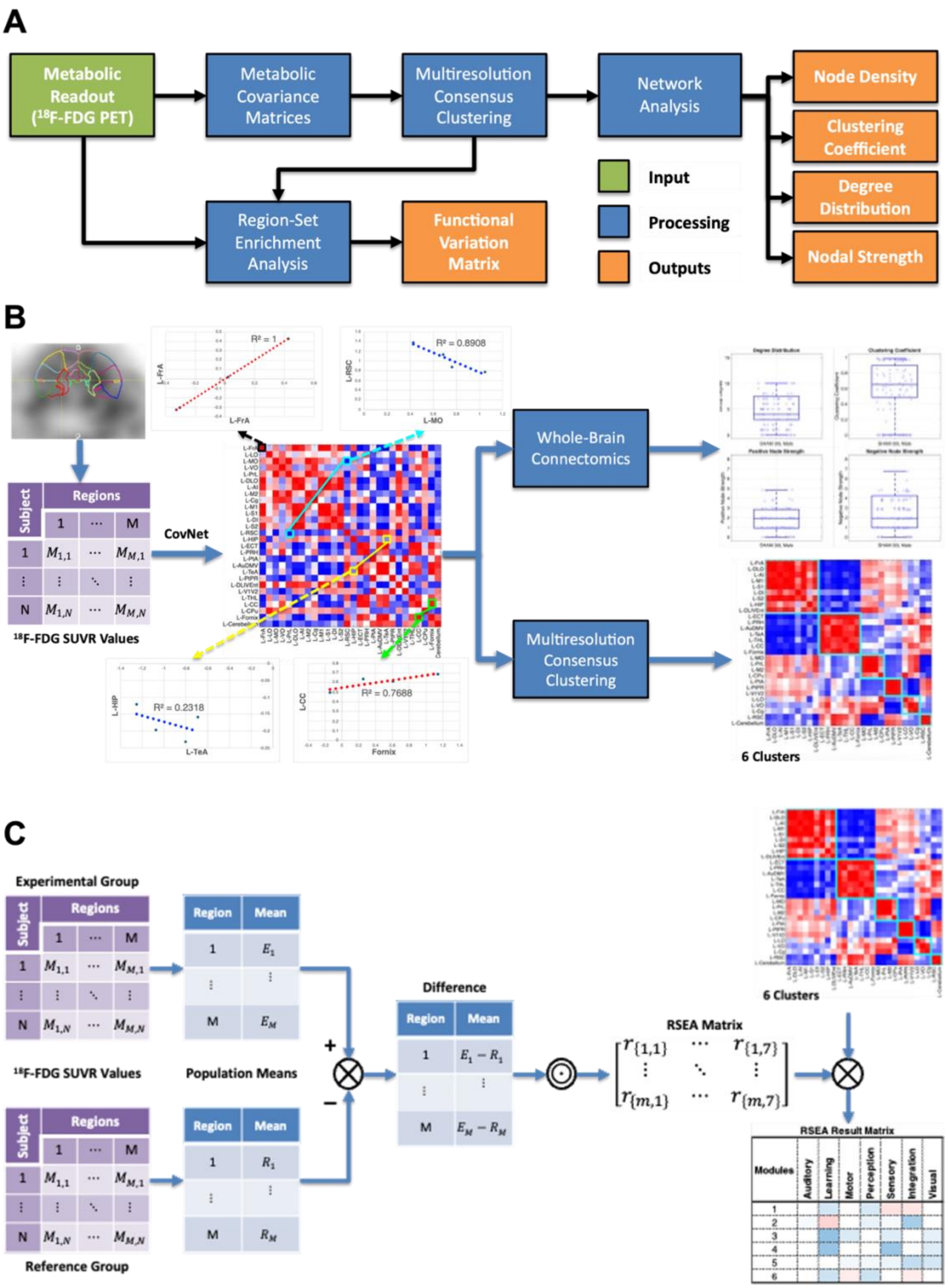

**Supplemental Figure S2 – Analysis Workflows for Metabolic Connectomics and Region-Set Enrichment Analysis (RSEA).**
**(A)** Schematic representation of the metabolic functional connectomics (MFC) analysis workflow used to evaluate network organization and changes in metabolic complexity at both whole-brain and regional levels. **(B)** Detailed workflow for the whole-brain MFC analysis pipeline, illustrating the construction of the metabolic covariance matrices, brain regions clustering using multi-resolution consensus clustering (MRCC), and derivation of the network metrics. **(C)** Depiction of the Region-Set Enrichment Analysis (RSEA) workflow, showing how $^{18}$F-FDG imaging data is used to compute region-wise differences between two groups to generate an RSEA heatmap that quantitatively detects brain function changes.

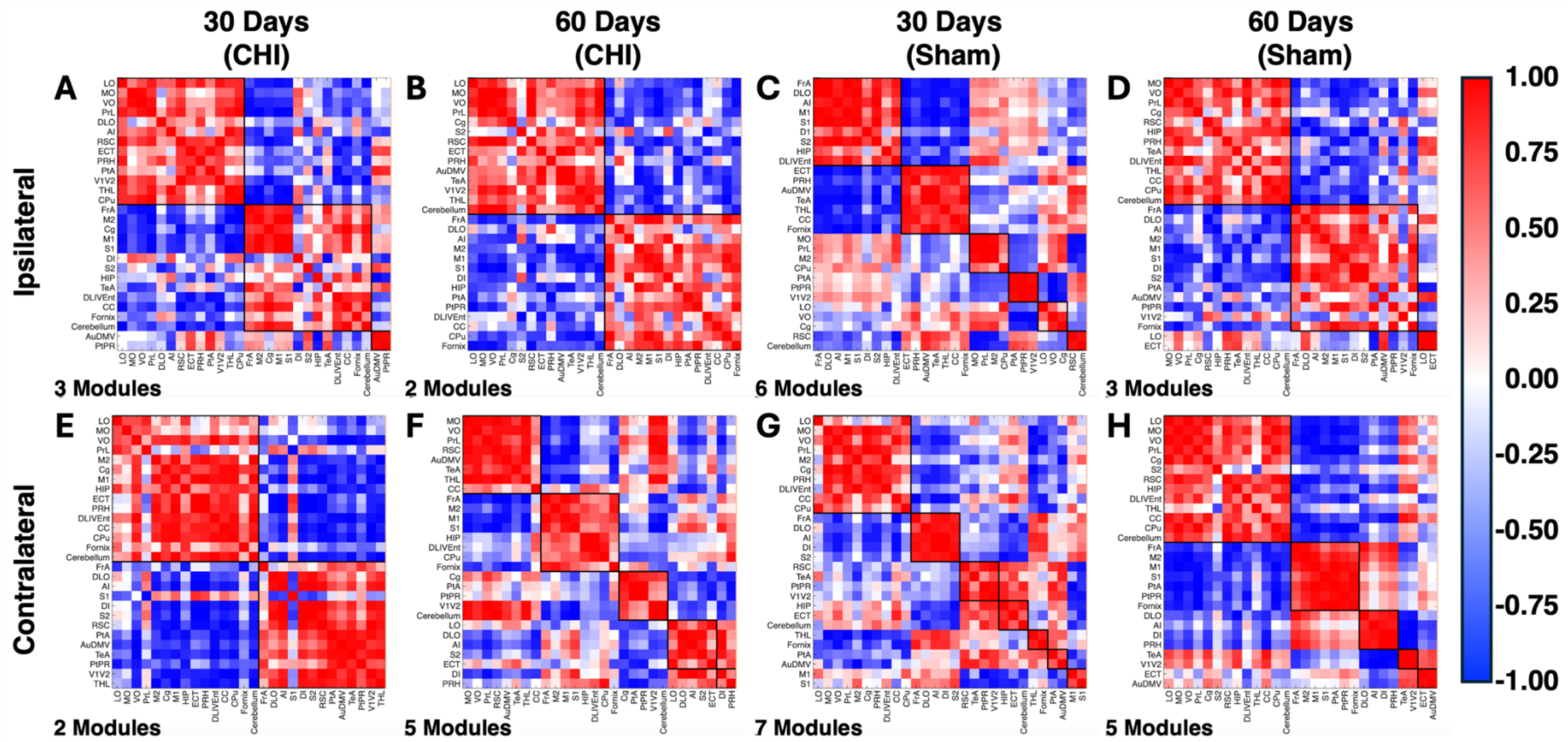

**Supplemental Figure S3 – Metabolic Covariance Matrices for Male Mice.**
Metabolic covariance matrices are shown for **(A-D)** ipsilateral (ILH) and **(E-H)** contralateral (CLH) hemispheres for **(A, E)** CHI at 30-DPI, **(B, F)** CHI at 60-DPI, **(C, G)** Sham at 30-DPI, and **(D, H)** Sham at 60-DPI. These matrices are symmetric, where each element is the pairwise Pearson correlation coefficient between brain regions, with warmer colors indicating positive correlations and cooler colors indicating negative correlations. Black outlines denote the regional MFC modules (RMCMs) identified using multi-resolution consensus clustering (MRCC) for each group. Full names of region labels are available in **Table S1**. Across hemispheres and time points, both the number and composition of RMCMs vary, indicating that brain regions dynamically reassign between modules. This metabolic network reorganization may be driven by both injury and aging.

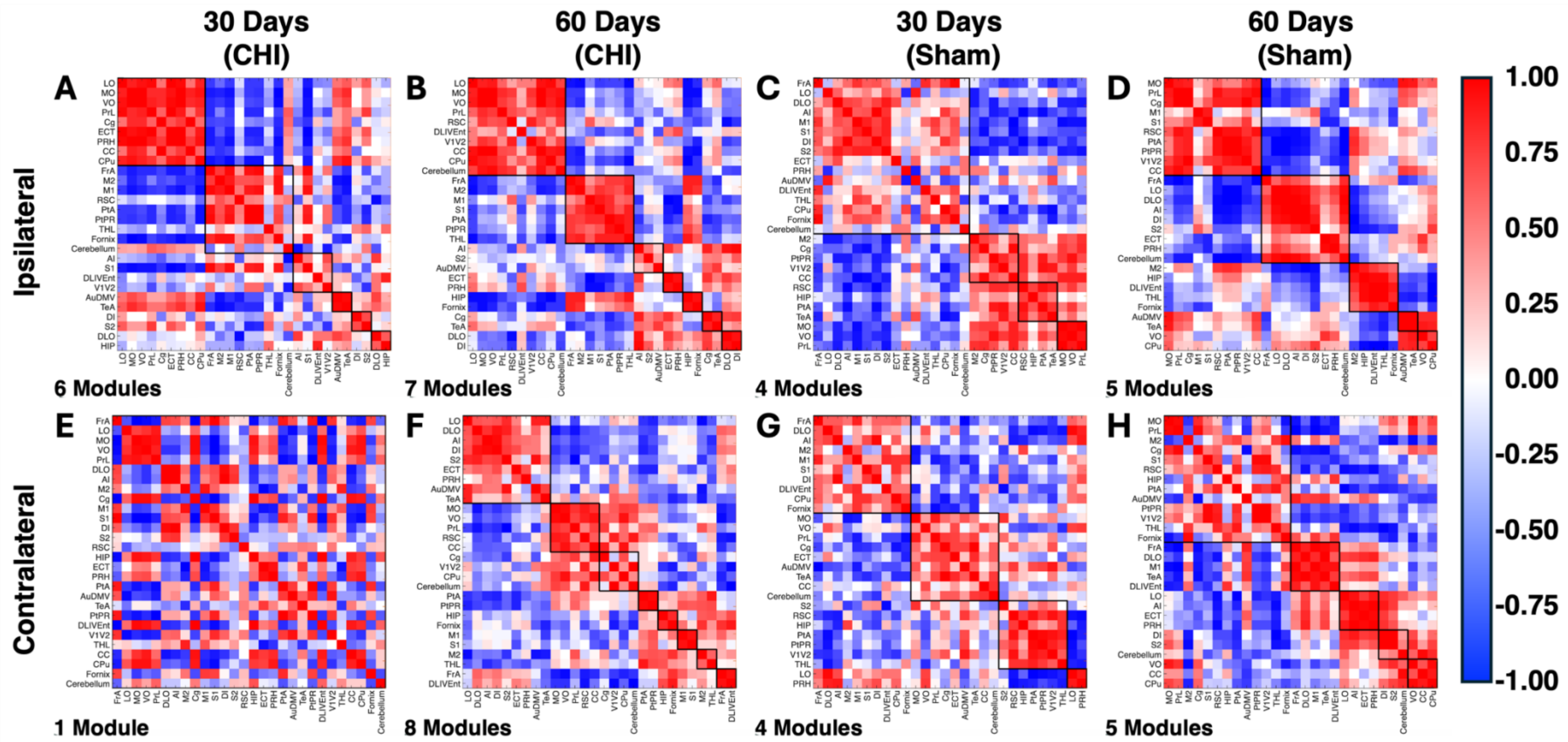

**Supplemental Figure S4 – Metabolic Covariance Matrices for Female Mice.**
Metabolic covariance matrices are shown for **(A-D)** ipsilateral (ILH) and **(E-H)** contralateral (CLH) hemispheres for **(A, E)** CHI at 30-DPI, **(B, F)** CHI at 60-DPI, **(C, G)** Sham at 30-DPI, and **(D, H)** Sham at 60-DPI. These matrices are symmetric, where each element is the pairwise Pearson correlation coefficient between brain regions, with warmer colors indicating positive correlations and cooler colors indicating negative correlations. Black outlines denote the regional MFC modules (RMCMs) identified using multi-resolution consensus clustering (MRCC) for each group. Full names of region labels are available in **Table S1**. As observed in males, the composition of RMCMs varies across groups. Notably, unlike males, the number of RMCMs did not vary between hemispheres in Sham at both time points.

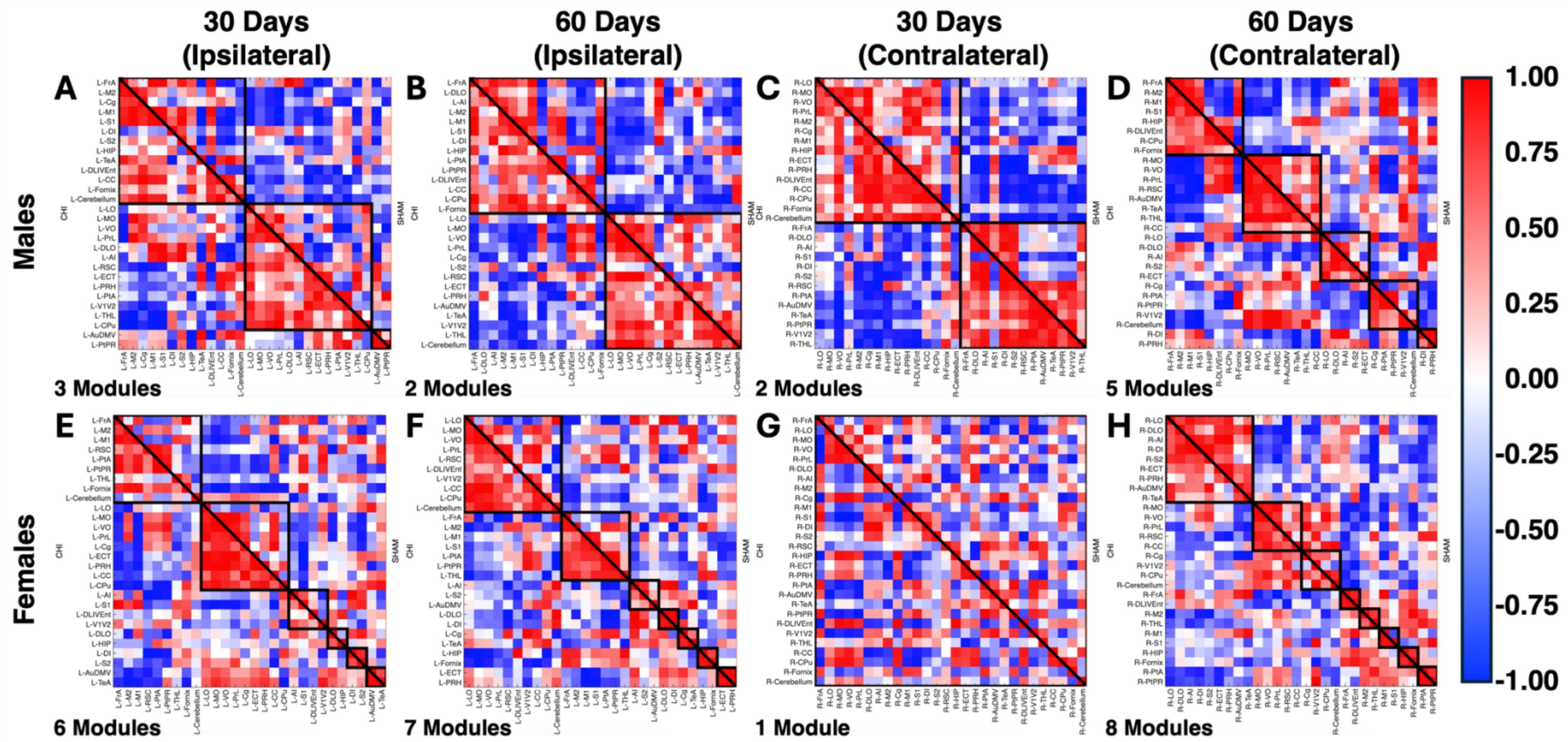

**Supplemental Figure S5 – Regional-Brain Analysis of Sham Relative to CHI.**
Regional metabolic functional connectomics (MFC) analysis using metabolic covariance matrices for **(A-D)** males and **(E-H)** females for **(A, E)** ILH at 30-DPI, **(B, F)** ILH at 60-DPI, **(C, G)** CLH at 30-DPI, and **(D, H)** CLH at 60-DPI. The upper triangular matrix is the comparison group (Sham), and the lower triangular matrix is the reference (CHI). Black outlines denote regional MFC modules (RMCM) identified using multi-resolution consensus clustering (MRCC) for CHI at 30-DPI. Data with warmer colors show positive correlations, while cooler colors show negative correlations. When CHI is treated as the reference group, Sham networks appear disorganized at 30-DPI, which may be due to CHI-derived RMCMs promoting changes in brain region clustering as a potential compensatory mechanism under injury conditions. However, as exhibited in the CHI relative to Sham comparison, metabolic covariance matrices in CHI and Sham mice begin to resemble one another at 60-DPI.

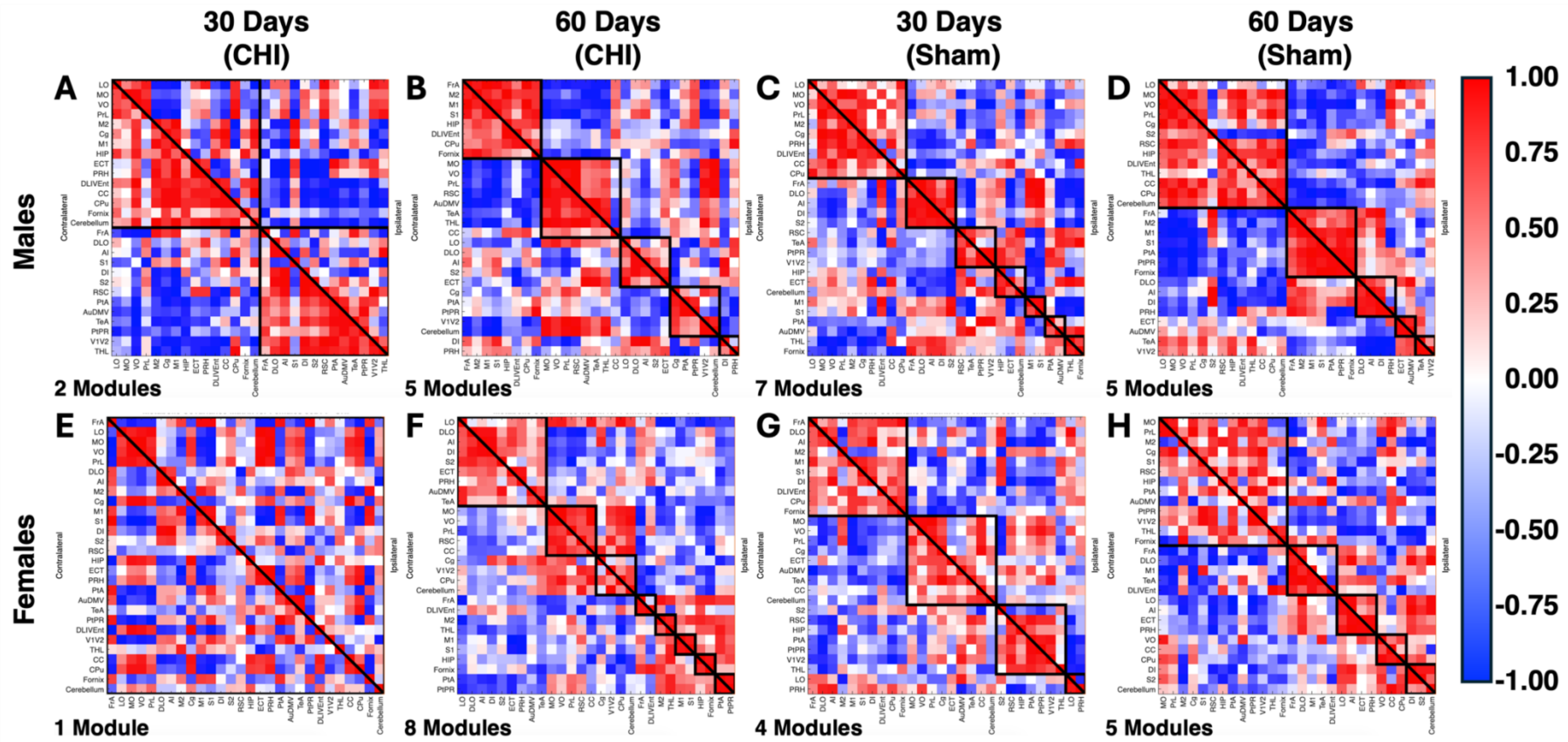

**Supplemental Figure S6 – Regional-Brain Analysis of Ipsilateral (ILH) Relative to Contralateral (CLH).** Regional metabolic functional connectomics (MFC) analysis using metabolic covariance matrices for **(A-D)** males and **(E-H)** females for **(A, E)** CHI at 30-DPI, **(B, F)** CHI at 60-DPI, **(C, G)** Sham at 30-DPI, and **(D, H)** Sham at 60-DPI. The upper triangular matrix is the comparison group (ILH), and the lower triangular matrix is the reference (CLH). Black outlines denote regional MFC modules (RMCM) identified using multi-resolution consensus clustering (MRCC) for CLH at 30-DPI. Data with warmer colors show positive correlations, while cooler colors show negative correlations. In CHI mice, inter-hemispheric comparisons (ILH relative to CLH) revealed a similar trend to the previous comparisons. At 30-DPI, both hemispheres show disorganization, which becomes less apparent at 60-DPI. Moreover, these findings suggest that both hemispheres in CHI mice may function similarly at 60-DPI. In Sham mice, differences between the hemispheres are minimal, as they show similar trends at both time points, consistent with the absence of injury.

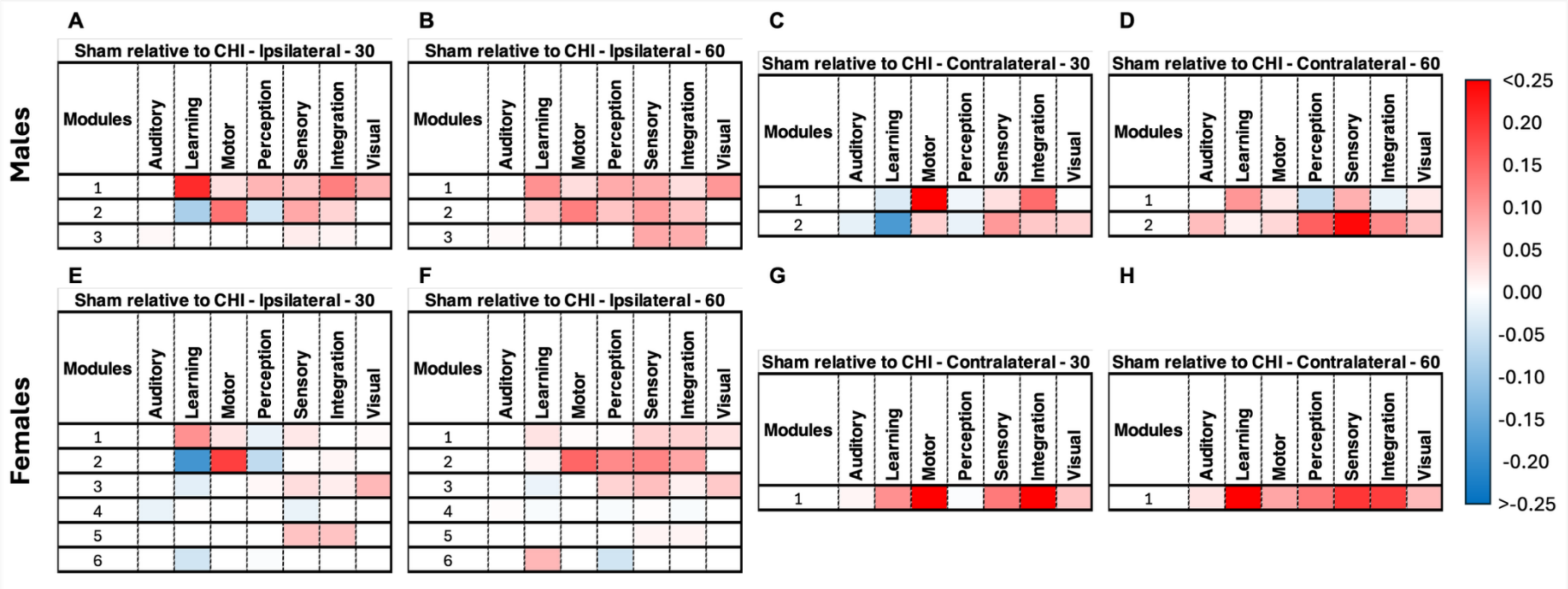

**Supplemental Figure S7 – RSEA of Sham Relative to CHI using 30-DPI CHI RMCM.**

Region-set enrichment analysis (RSEA) heatmaps are shown for **(A-D)** males and **(E-H)** females. These are split as follows: **(A, E)** ILH at 30-DPI, **(B, F)** ILH at 60-DPI, **(C, G)** CLH at 30-DPI, and **(D, H)** CLH at 60-DPI. RMCM were obtained using MRCC on the Sham mice at 30-DPI for each hemisphere. These heatmaps show metabolic changes across seven functional categories, based on $^{18}$F-FDG readouts. Data with warmer colors shows positive fractional percentage changes, while cooler colors show negative fractional percentage changes. In these RSEA heatmaps, positive changes indicate a fractional percentage increase in metabolic demand for a given functional category in the comparison group (CHI) relative to the reference (Sham), while negative changes indicate a relative decrease, and values near 0 indicate similarity between groups. When comparing Sham relative to CHI, Sham mice of both sexes exhibit higher metabolic changes than CHI mice across both time points and hemispheres, except within the auditory category. These findings suggest that CHI mice continue to exhibit reduced metabolic activity, supporting the idea that CHI-induced injury dampens overall brain network metabolic complexity.

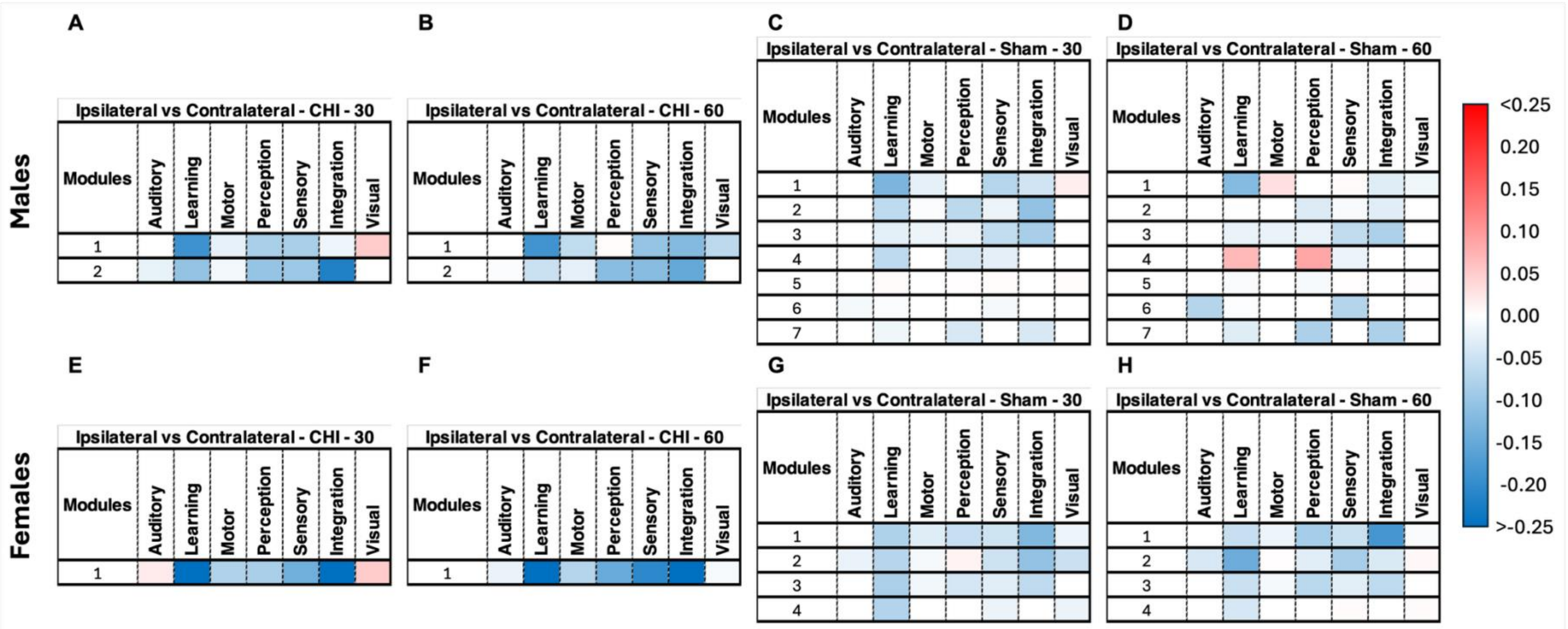

**Supplemental Figure S8 – RSEA Hemispheric Comparison (Ipsilateral (ILH) Relative to Contralateral (CLH)) using 30-DPI CLH RMCM.** Region-set enrichment analysis (RSEA) heatmaps are shown for **(A-D)** males and **(E-H)** females. These are split as follows: **(A, E)** ILH at 30-DPI, **(B, F)** ILH at 60-DPI, **(C, G)** CLH at 30-DPI, and **(D, H)** CLH at 60-DPI. RMCM were obtained using MRCC on the Sham mice at 30-DPI for each hemisphere. These heatmaps show metabolic changes across seven functional categories, based on $^{18}$F-FDG readouts. Data with warmer colors shows positive fractional percentage changes, while cooler colors show negative fractional percentage changes. In these RSEA heatmaps, positive changes indicate a fractional percentage increase in metabolic demand for a given functional category in the comparison group (CHI) relative to the reference (Sham), while negative changes indicate a relative decrease, and values near 0 indicate similarity between groups. In CHI mice of both sexes, inter-hemispheric comparisons reveal overall decreases in ILH metabolic changes across most functional categories at 30-DPI, with only a subtle attenuation at 60-DPI. These findings indicate that CHI injuries result in a persistent deficit in ILH relative to CLH, which is not recovered by 60-DPI. In contrast, Sham mice exhibit minimal hemispheric differences at either time point.
